# More than 100 dual coding regions have evidence for selection constraints in both reading frames

**DOI:** 10.64898/2026.05.28.727796

**Authors:** Michael L. Tress, Daniel Cerdán-Vélez, Miguel Maquedano, Federico Abascal

## Abstract

Alternative splicing can generate multiple differently spliced transcripts from a single pre-mRNA. A striking number of genes have alternative splice events that ccan hange the downstream reading frame leading to exons that code from distinct reading frames. In fact, more than a third of the coding genes in the human gene set are annotated with dual coding exons derived from alternative splicing events.

Here we analysed a set of 537 dual coding regions that have evidence to support their functional importance. These dual coding regions produce protein isoforms with completely different C-terminals and have reading frames that are supported by either peptide or conservation evidence. More than a quarter of the alternative reading frames are preserved across all mammals, and many can be traced back to the earliest jawed vertebrates. Most of these ancient dual coding regions appear to be under selective constraints. We find support for purifying selection on both frames in 105 pairs of transcripts and two genes, *CCSER2* and *SH2B1*, have triple coding regions that are under clear selection pressure in all three frames.

We found evidence to suggest that many ancient dual coding regions may have played important roles in the evolution of the vertebrate central nervous system. Most ancient dual coding regions with evidence for protein level tissue specificity were brain specific and we showed that genes with ancient dual coding regions are highly enriched in brain tissues. Most remarkably, we found that more than 80% of the genes with these ancient dual coding regions are implicated in neuron development, synapses and neural cell projections.

## Introduction

Almost all multi-exon protein coding genes can generate alternatively spliced transcripts (Wang *et al*, 2008) and with the advent of long read transcriptomics analyses the number of differently spliced transcripts per gene with confirmatory evidence has exploded (Perteghella *et al*, 2026). Alternative splicing is one potential mechanism for introducing novel protein functions in multicellular organisms (Merkin *et al*, 2012; Vuong *et al*, 2016; Wright *et al*, 2022). However, the extent to which alternatively spliced protein isoforms contribute to cellular function is not clear beyond a few well studied examples (Wright *et al*, 2022).

Most coding genes have a single main protein isoform (Ezkurdia *et al*, 2015, Pozo, Martinez Gomez *et al*, 2022, Pozo, Rodriguez *et al*, 2022). By way of contrast, a large majority of alternative transcripts are not under selective pressure (Liu and Lin, 2015; Tress *et al*, 2017; Pozo, Martinez Gomez, *et al*, 2022), proteomics analyses detect relatively little evidence for their translation (Ezkurdia *et al*, 2015; Rodriguez *et al*, 2020) and very few capture validated ClinVar (Landrum *et al*, 2018) pathogenic mutations (Pozo, Rodriguez, *et al*, 2022).

A minority of alternative protein isoforms appear to be functionally important (Tress *et al*, 2017), and these isoforms are almost always of ancient origin (Martinez Gomez *et al*, 2020, Rodriguez *et al*, 2020). Many of these conserved alternative isoforms are tissue specific (Rodriguez *et al*, 2020) and they do have associated validated pathogenic mutations (Pozo, Rodriguez, *et al*, 2022). One class of splice isoforms stands out in this regard, those generated from tandem duplicated exon substitutions (Martinez Gomez *et al*, 2020, Martinez Gomez *et al*, 2021).

Classical alternative splicing envisages that alternative transcripts are formed from the addition or deletion of exonic sequences (Wright *et al*, 2022). If the exonic sequence that is gained or lost is a multiple of three, the alternative transcript will produce a protein isoform that is identical save for a region of missing or added amino acids. If it is not, downstream exons will be read in a different reading frame, and the corresponding transcripts will produce protein isoforms with radically different C-termini.

Exons that can code from two different reading frames (dual coding regions) are particularly interesting because the exon must be under special evolutionary constraints if both reading frames are functionally important. Since almost all mutations that are synonymous in one frame are non-synonymous in the other, practically all nucleotide changes will lead to a change of amino acid sequence in one protein or the other (Chung *et al*, 2007).

Dual coding regions can be generated via a range of mechanisms, alternative splicing, out of frame translation initiation sites, adjacent overlapping coding genes and programmed ribosomal frameshifting. Dual coding regions that stem from distinct upstream translation initiation sites have received a lot of attention, and there are several well-known examples, such as the ARF and p16-INK4a proteins in *CDKN2A* (Szklarczyk *et al*, 2007). These dual coding exons, and those produced from adjacent genes that overlap such as *MOCS2* (Reiss, 2000), produce entirely different proteins that (if functional) will have different roles from the principal isoforms. Programmed ribosome frameshifting (Cagliani *et al*, 2024) is much more common in bacteria and viruses, but among human coding genes there are clear examples such as *PLEKHM2* (Loughran *et al*, 2025) and the retroviral-derived *PEG10* (Shigemoto *et al*, 2001).

Alternative splicing derived dual coding regions events are by far the most common. More than a third of human coding genes (6,835 coding genes in GENCODE v45, Mudge *et al*, 2025) are annotated with at least one transcript with an alternative splicing derived dual coding region. These dual coding regions usually generate alternative transcripts with premature stop codons and will therefore produce alternative proteins with distinct C-terminal regions that are shorter than the principal isoform.

*A priori*, it might be imagined that frame changing splice events would be less likely to produce functional proteins, since changing the reading frame abolishes any functional and structural domains and motifs downstream of the splice event and replaces them with quasi random amino acid sequences. In addition, evolutionary adaptation would be limited by the selection constraints that already exist on the principal coding frame.

There have been three genome-wide analyses of dual coding regions. Liang and Landweber (Liang and Landweber, 2006) found 173 genes annotated with dual coding frames (and another three with triple coding frames). They found that most alternative reading frames evolved recently and predicted that most, if not all, of the alternative transcripts were functional. Kovacs *et al* (Kovacs *et al*, 2010) analysed 97 overlapping reading frames that were longer than 75 nucleotides and that had unspecified protein evidence. They found that alternative frames had more residues typical of disordered regions than the principal frames, which would mean that they were less likely to be degraded by cellular surveillance mechanisms (Casola *et al*, 2026).

The third study (Goubert *et al*, 2026) concentrated on the almost 1300 dual coding regions that are annotated in the UniProtKB human proteome entries. The study confirmed that UniProtKB dual coding regions were generated overwhelmingly from alternative splicing and would produce proteins with C-terminals that are unlikely to fold into globular structures. They also predicted that many UniProtKB dual coding regions were likely to be involved in gene regulation since the majority were probably degraded via the nonsense mediated decay pathway.

The dual coding regions annotated by UniProtKB and GENCODE are both manually curated. There are more than ten times as many dual coding regions in GENCODE transcripts as there are in UniProtKB proteins. Another advantage is that all dual coding regions map to the human genome assembly, something that is not always true of UniProtKB manually curated isoforms (Maquedano *et al*, 2025). However, the GENCODE human gene set is annotated with tens of thousands of transcripts with dual coding regions, and it is difficult to know which are most likely to be functionally relevant.

Here, we investigated whether there is any evidence to support functional roles for alternative protein isoforms produced from dual coding regions. We started with a set of 537 dual coding regions that either had clear cross-species conservation evidence or had peptide evidence supporting translation from both frames. Conservation is a clear marker of biological relevance, and we used peptide evidence as a proxy for biological function because isoforms with proteomics support must be more highly expressed and must have avoided cellular surveillance mechanisms (Kesner *et al*, 2023; Tress, 2025). We found that more than a hundred conserved dual coding regions appear to be under selection pressure. Remarkably almost all the genes with ancient dual coding regions have roles in neuron development.

## Materials and Methods

### Proteomics analysis

For the proteomics analysis, we mapped validated human proteome peptides from the PeptideAtlas database (Desiere *et al*, 2006, January 2025 build) to proteins from the GENCODE v45 gene set (Mudge *et al*, 2025) from which the readthrough genes had been eliminated (Maquedano *et al*, 2025). We allowed only fully tryptic peptides, but missed tryptic cleavages were permitted. PeptideAtlas peptides had a minimum of 7 amino acids and a maximum of 50 amino acids. Peptides that mapped to more than one gene were not considered.

In order to identify a dual coding region, we required that tryptic peptides mapped to both reading frames of the dual coding exons. Valid peptides could either map to the dual coding region itself or to the dual coding exons outside of the dual coding region (see figure 1). We identified a dual coding region with peptides if both frames had a minimum of two supporting PeptideAtlas observations. PeptideAtlas peptides supported 430 dual coding regions in 412 distinct GENCODE v45 genes.

**Figure 1.**
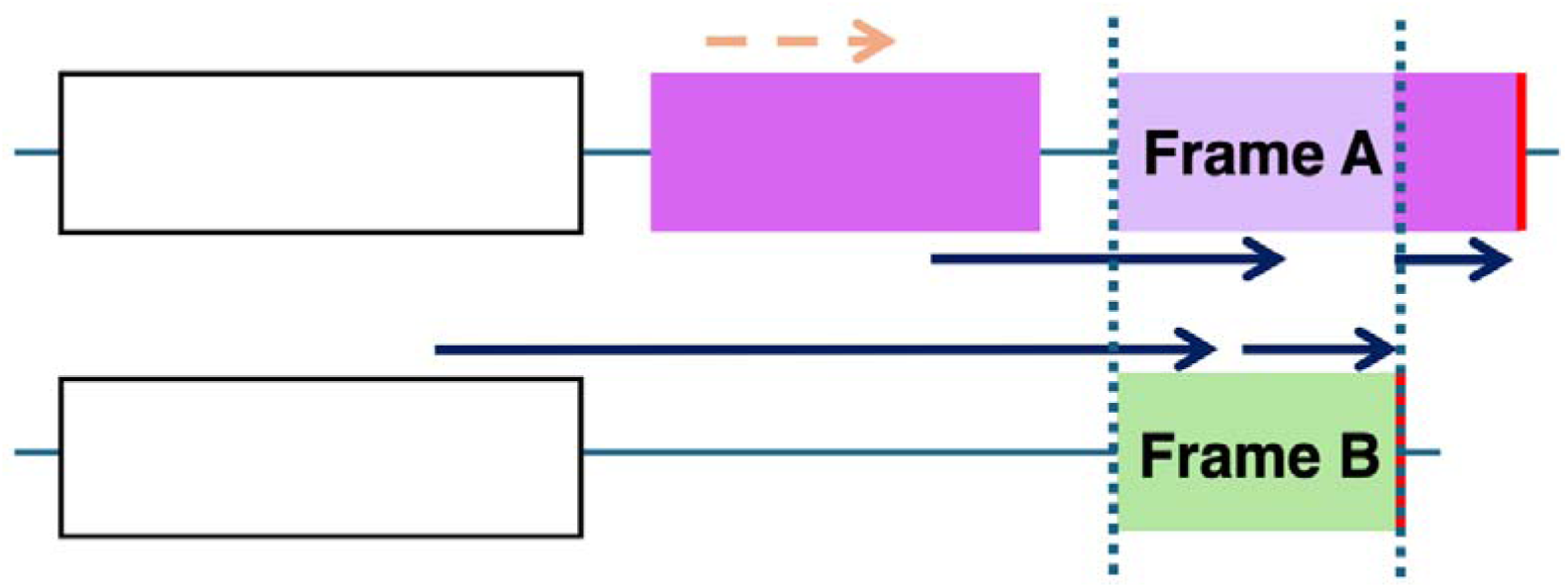
Peptide validation of dual coding regions. A schematic representations of two transcripts with dual coding regions at the 3’ ends illustrating their validation with PeptideAtlas peptides. Exons in different colours are in different coding frames, the two coding frames (caused in this case by an exon skip) are shown in green and purple. The region of the dual coding exons with overlapping reading frames is marked by colours with a lighter shade and bounded by vertical dashed lines. The stop codons are shown as red vertical lines. Valid peptides (that overlap the dual coding exon) are shown as black arrows. The dashed orange arrow shows the position of an invalid peptide (although it may be unique to the transcript, it does not overlap the dual coding exon). Both reading frames had to be supported by valid peptides

### Manual detection of dual coding regions with peptide or conservation support

In addition to the 430 dual coding regions annotated in the Ensembl/GENCODE gene set that had peptide support for both reading frames, we found another 107 dual coding regions via manual curation. These dual coding regions either had peptide or conservation evidence and came from two sources. The first source was a published analysis in which we had used the PeptideAtlas database to search for coding regions missing from the GENCODE v45 gene set (Rodriguez *et al*, 2025). By mapping PeptideAtlas peptides that did not match GENCODE v45 isoforms back to the genome, we had found support for 279 alternative splicing events that were not annotated in the Ensembl/GENCODE gene set (Rodriguez *et al*, 2025). Included among these splice events were 65 dual coding 3’ exons. Of these dual coding regions, 49 were annotated in RefSeq (Goldfarb *et al*, 2025), a further 8 were annotated as part of proteins in UniProtKB (UniProtKB, 2025) and 8 were unannotated predictions from large-scale experiments (Deutsch *et al*, 2016).

The second source came from an assumption that genes with ancient dual coding regions might have paralogues with equivalent dual coding regions. This search turned up another 42 dual coding regions that had highly conserved dual-coding reading frames. These dual coding regions did not have peptide support, 29 of these dual coding regions were annotated in either the RefSeq or Ensembl/GENCODE reference gene sets, but 13 were not annotated as part of coding transcripts.

### Determining main isoforms

For each dual coding region, we chose a pair of transcript and isoforms (one for each coding frame) and labelled them as main and alternative for technical reasons. We have shown that protein isoforms coincide overwhelmingly (Pozo, Martinez Gomez *et al*, 2022, Pozo, Rodriguez *et al*, 2022) with APPRIS principal isoforms (Rodriguez *et al*, 2022), so in those cases where one of the transcripts coincided with a principal transcript, we chose it as the main transcript. if only one of the two transcripts was annotated in Ensembl/GENCODE, we chose the Ensembl/GENCODE transcript as the main transcript. For the remaining transcripts we chose the isoform that had more peptide evidence as the main isoform.

The main transcript almost always included the reading frame with the most cross species conservation, though this was not always the case.

### Preservation of reading frames

We determined the depth of the cross-species evidence for the reading frames in each dual coding region based on alignments from the UCSC Genome Browser (**Perez *et al*, 2025**) that are available in the codalignview web pages (https://data.broadinstitute.org/compbio1/cav.php). We analysed the alignments for each predicted reading frame in the dual coding region, not just over the dual coding region of each dual coding exon, but also over each dual coding exon from the 5’ end of the exon to the stop codon or the 3’ end of the exon (whichever came first). We always required both reading frames to be preserved.

We used three different sets of alignments to determine whether a reading frame was preserved across species, the 447-way mammalian alignments (Kuderna *et al*, 2024), the 470-way mammalian alignments (Raney *et al*, 2024) and the 100 vertebrate alignments (Rosenbloom *et al*, 2015). While the 100-species vertebrate alignments had the furthest reach (including multiple bird and reptile species), there are few primate species, and the alignments are older and less reliable. The 470-way mammalian alignments had the most depth and were the most recent alignments, but the 447-way alignments had the largest range of primate species. If there were disagreements between any of the three sets of alignments, we used the results from the 470-way alignments because they were the most recent.

A reading frame was determined to be preserved across a clade if there were no (or almost no, see below) premature stops or frameshifts in the reading frame of the orthologues, there were no lost stop codons in the orthologues and any frame-changing splice sites were also conserved across orthologues.

Given that there may be sequencing errors in some of the species, we tolerated one or two frame disrupting changes in a clade, as long as the frame was preserved in more distant clades. We were particularly tolerant of frameshifts in the dual coding region, assuming that these were errors, since these would affect both frames. We did find a small number of cases in which a reading frame was lost in one family, but was clearly maintained beyond that, as was the case with one of the two dual coding reading frames in *CCSER2*. This reading frame had premature stop codons in bats but was clearly maintained across all jawed vertebrates apart from that.

The “age” of the dual coding frame was determined to be the most distant clade in which both frames of the exon or exons were completely preserved. On its own, frame preservation is one indication that a dual coding region may have gained a functional role, but it is not sufficient since alternative reading frames that overlap completely with the reading frames of the principal transcript (see figure 1) may only have preserved reading frames due to selection pressures within the main reading frame. This is particularly true if the dual coding region is short.

The cross-species alignments served to indicate whether reading frames were preserved across mammals, amniotes and tetrapods, though the 100-species vertebrate alignments only include one amphibian species, *Xenopus tropicalis*. To determine whether alternative reading frames and stop codons were preserved beyond amniotes, we looked for evidence of annotated proteins with BLAST searches against more distant clades and carried out direct analysis of the equivalent exons in fish, elephant shark, lamprey and lancelet genomes in the UCSC browser.

### Tissue Specificity

For the 495 dual coding regions that had PeptideAtlas peptides for both reading frames (both the 430 dual coding regions annotated in GENCODE v45 and the 65 that were not annotated), we used the peptides and the tissues in which they were detected to determine whether there was any evidence for tissue specificity at the protein level. The 3,104 experiments that made up PeptideAtlas were a mixture of diseased and cancerous tissues, cell lines and healthy tissues. For this analysis, we discarded the cancer and disease observations as well as those from cell lines and concentrated just on experiments that were carried out on healthy tissue.

Almost all PeptideAtlas analyses have an associated name, which clearly denotes the tissue or cell line. When there were doubts about the tissue used, there was always further information from paper abstracts that detailed whether the investigated tissue was diseased or healthy. We pooled the tissues into broad categories including nervous tissue, heart/skeletal muscle, blood/immune system, digestive system, to boost the number of experiments in each category.

Proteomics experiments, particularly large-scale experiments, which many of the PeptideAtlas analyses were, often have different ways of dividing their analysis into separate individual experiments. In PeptideAtlas each experiment has an identity number, so to simplify matters, we used this number to identify individual experiments. For each of the tryptic peptides that uniquely identified a reading frame, we simply counted the number of individual tissue-based experiments that the peptide was observed in. For each group of tissues, we summed the number of times peptides that mapped to each of the two reading frames were identified in PeptideAtlas experiments.

To determine whether isoforms coded from the two frames were significantly enriched in a specific tissue, we used Fisher’s exact tests based on the total peptide identifications for each tissue group and for each reading frame.

### Gene expression

Transcript expression for Ensembl/GENCODE genes is available from the Human Protein Atlas (Uhlen *et al*, 2015). There were three tissue-based expression analyses available for Ensembl/GENCODE coding genes, an expression analysis from the Human Protein Atlas consortium itself (Uhlen *et al*, 2015), which has transcript expression levels for 40 distinct tissues, the GTEx analysis (GTEx Consortium, 2020) with expression levels consolidated into 33 tissues, and the FANTOM analysis (Abugessaisa *et al*, 2017), which has been consolidated into 46 tissues.

For each gene we normalised the tissue expression against the tissue with the highest expression. To calculate the mean expression over different age ranges, we binned the 412 genes with GENCODE v45 dual coding regions supported by PeptideAtlas peptides by the predicted age of their dual coding regions. Clades with few genes were grouped with adjacent clades.

### Strategies to detect evolutionary constraints

In all but a handful of dual coding regions, the selection pressure on one reading frame is stronger than the selection pressure in the other. In those cases where the selection pressure appears to be more or less equal for both frames, such as the gene *SYNCRIP*, the overlapping region is so conserved that it is not possible to measure selective pressure in either frame. Even if both overlapping frames are under selection, estimation of dN/dS where the two frames overlap region is complicated because strong purifying selection on one frame influences selection on the other. However, three strategies that allowed us to assess the evidence for selection pressure on both reading frames of a dual coding region. These strategies are detailed here.

Firstly, if one of the two reading frames extends sufficiently beyond the other, we were sometimes able to measure dN/dS for two distinct frames using the same exon. This applied only when the shorter reading frame was the one that was under stronger selection pressure (Figure 2A). In these cases, we carried out dN/dS analyses using the overlapping section of the dual coding exon for the frame that was under stronger selection pressure (frame B in Figure 2A) and the region that extended beyond the dual coding section for the frame that was under weaker selection pressure (frame A in Figure 2A). For the dN/dS analysis both the overlapping section of the dual coding region and the section that extended beyond the overlapping section had to be at least 3 codons in length.

**Figure 2.**
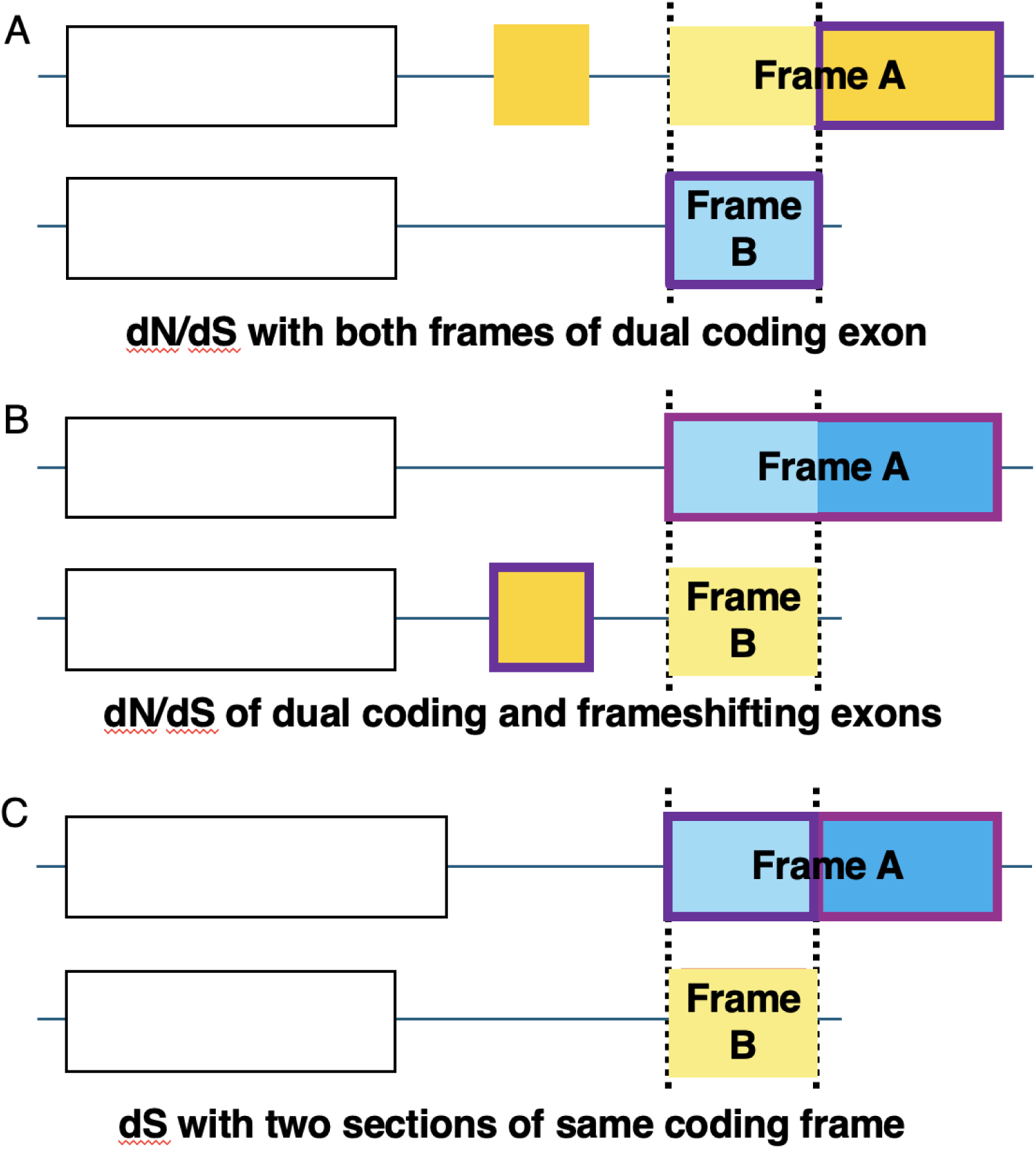
Testing selection in dual coding regions. Schematic representations of dual coding regions illustrating the three strategies for selection testing. Exons in white are common to both transcripts. Exons in different colours are in different coding frames, the overlapping regions of the dual coding exons are marked by colours with a lighter shade and bounded by vertical dashed lines. In all three panels the reading frame in blue is under stronger selection pressure than the yellow reading frame. The two purple boxes in each panel show the two regions that are tested. A. The reading frame under weaker selection pressure extends beyond the overlapping section, so we can calculate dN/dS for two different sections of the dual coding exon. B. We cannot calculate dN/dS for the weaker yellow frame because the blue coding frame fully overlaps it, instead we calculate the dN/dS of the exon (or exon extension) that causes the change of frame. C. We cannot calculate dN/dS for the weaker yellow frame and there is no frame-changing exon, but we can calculate dS over two sections of the blue reading frame, for the overlapping section of dual coding exon (light blue) and for section that codes from just one reading frame that extends beyond the overlapping frames (darker blue).

If the longer of the two overlapping coding frames was the frame under stronger selection constraints (for example, in Figure 2B and 2C), the first strategy was not practicable. In these cases, a second strategy could be applied if the shorter reading frame was generated from an inserted exon or exon extension (Figure 2B). This allowed us to estimate whether the alternative frame was under selection pressure by calculating the dN/dS of the frame changing exon rather than the dual coding exon itself. We only carried out this analysis if the alternative reading frame was associated uniquely with the upstream frame changing exon/extension. For example, we could not validate selection for the dual coding exons in the ATPase 1 family via this method because although the frame changing exons were under selection pressure, these exons were also present in transcripts other than those with the dual coding regions. For several dual coding regions, we joined the frame changing exon to the overlapping frame of the dual coding exon to measure dN/dS ratios.

Where the longer of the two frames in the dual coding region was under greater selection constraints and there was no frame-changing exon, a third strategy allowed us to estimate selection pressure. In this strategy, dS rates were calculated for the dual coding region over the reading frame that was under stronger selection pressure (Figure 2C). We calculated dS values from two different sections of this principal reading frame, the section that was dual coding and the section that extended beyond the dual coding exon. If the dual coding region were under selection pressure in two different frames, we would expect to see a substantially lower dS rate for the dual coding section of the exon than for the extended section where selection pressure is only acting on one frame.

### Comparative dN/dS analysis

To calculate dN/dS ratios, we generated cross-species alignments for each dual coding region analysed. We used alignments either from the 470-way mammalian alignments, the 447-way mammalian alignments or the 100-way vertebrate alignments, based on the predicted age of the frame. For example, if both frames were only preserved across placental mammals, we only included placental mammal species in the alignments that we used to calculate the dN/dS. We removed all sequences that had frameshifts from the alignments along with those that did not exclusively align bases (for example that used “X” or “N”).

We estimated the maximum likelihood (ML) phylogenetic tree from each alignment using Phyml v3.3 (Guindon *et al*, 2010). The resulting tree was used to estimate global dN/dS ratios using Paml’s codeml program (Yang, 2007). We ran codeml twice for each alignment, one fixing dN/dS at 1, one with dN/dS as a free parameter. A likelihood ratio test (LTR) between the two models provides a p-value that indicates whether departure from neutrality is statistically supported. In summary, dN/dS significantly lower than 1 is evidence of purifying selection. Resulting p-values were adjusted for multiple hypothesis testing using Benjamini & Hochberg false discovery rate method (Benjamini and Hochberg, 1995).

### Comparative dS analysis

For the dS analysis of the dual coding regions, we created two cross species alignments. One covered the overlapping section of the exon based on the longer coding frame (frame A in figure 2C) and the other covered the section that extended beyond the dual coding region of the exon, based on the same longer coding frame. For dS calculations, both sections had to have a minimum of 5 codons. The idea here is: if the alternative frame is under selection, then we would see a reduction of synonymous substitutions in the current frame. We use the non-overlapping part of the exon as a reference of the background rate of synonymous substitutions for that gene and set of species.

As with the alignments for dN/dS analysis, we used alignments either from the 470-way mammalian alignments, the 447-way mammalian alignments or the 100-way vertebrate alignments, based on the predicted age of the frame. As with the dN/dS alignments, we removed all sequences that had frameshifts or that did not exclusively align bases. To be able to compare dS values between the two alignments, both had to have the exact same taxa, hence we removed any discrepancy.

As with the dN/dS analysis, we estimated ML phylogenetic trees with Phyml. Here we ran codeml once for each alignment with a free dN/dS parameter. The dS values (rate of synonymous substitutions per synonymous site) were obtained for each of the two alignments (overlapping and non-overlapping).

For the dS/dS analysis, we had to model the scores to be able to predict which dS/dS ratios were indicative of selection pressure in both frames. To model the dS/dS ratios, we took 20 dual coding regions that we knew were under selection pressure in both frames and 20 regions that we can be sure are not under selection pressure in both frames. For the first set we used those dual coding regions that we had already validated with dN/dS. For the second set, we used dual coding regions that were not preserved among simian species because their alternative frame had stop codons and/or lost stops.

## Results

### GENCODE dual coding regions with peptide support

We found PeptideAtlas peptide support for 430 GENCODE v45 alternative splicing-derived dual coding regions. These dual coding regions came from 412 different genes. The average length of the overlapping regions was 34.3 codons, while the longest dual coding region was in *LMTK3*, spanning 573 codons over 5 exons. From PeptideAtlas we calculated the number of peptides that mapped to each alternative reading frame and the number of times these peptides were observed in PeptideAtlas experiments (peptide observations). On average, there were 2.5 peptides per alternative reading frame. The *LMTK3* dual coding region was also the alternative reading frame with the most supporting peptides (49), while almost half of the 430 dual coding regions (199) had alternative frames that were supported by a single tryptic peptide. We required a minimum of 2 peptide observations as support for the translation of alternative frames and 42 of the 430 alternative frames were supported by the minimum 2 observations. At the other end of the scale, 76 of the dual coding regions had more than 100 observations, with a dual coding region in *HMGN3* having the most (4,346 observations).

### Estimating the age of dual coding regions

We estimated an age for each dual coding region based on its cross-species alignments. The estimated age was the age of the oldest clade in which both reading frames of the dual coding region were preserved. The process is detailed in the methods section. Dual coding regions were divided into 15 groups based on the oldest clade in which both frames were completely (or almost completely) conserved.

The distribution of the predicted ages of the dual coding regions with peptide support (see figure 3A), showed that the bulk of the dual coding reading frames with peptide support had little or no evidence of cross-species conservation and were recent evolutionary innovations at best. In 272 of the 430 dual coding frames with peptide evidence (63.3%) alternative reading frames were not even preserved across all primate orthologues. Although the majority of these dual coding regions did not show deep cross-species conservation, there were still at least 53 that were preserved beyond the ancestor of jawed vertebrates, more than 460 million years ago (Kumar *et al*, 2022).

**Figure 3.**
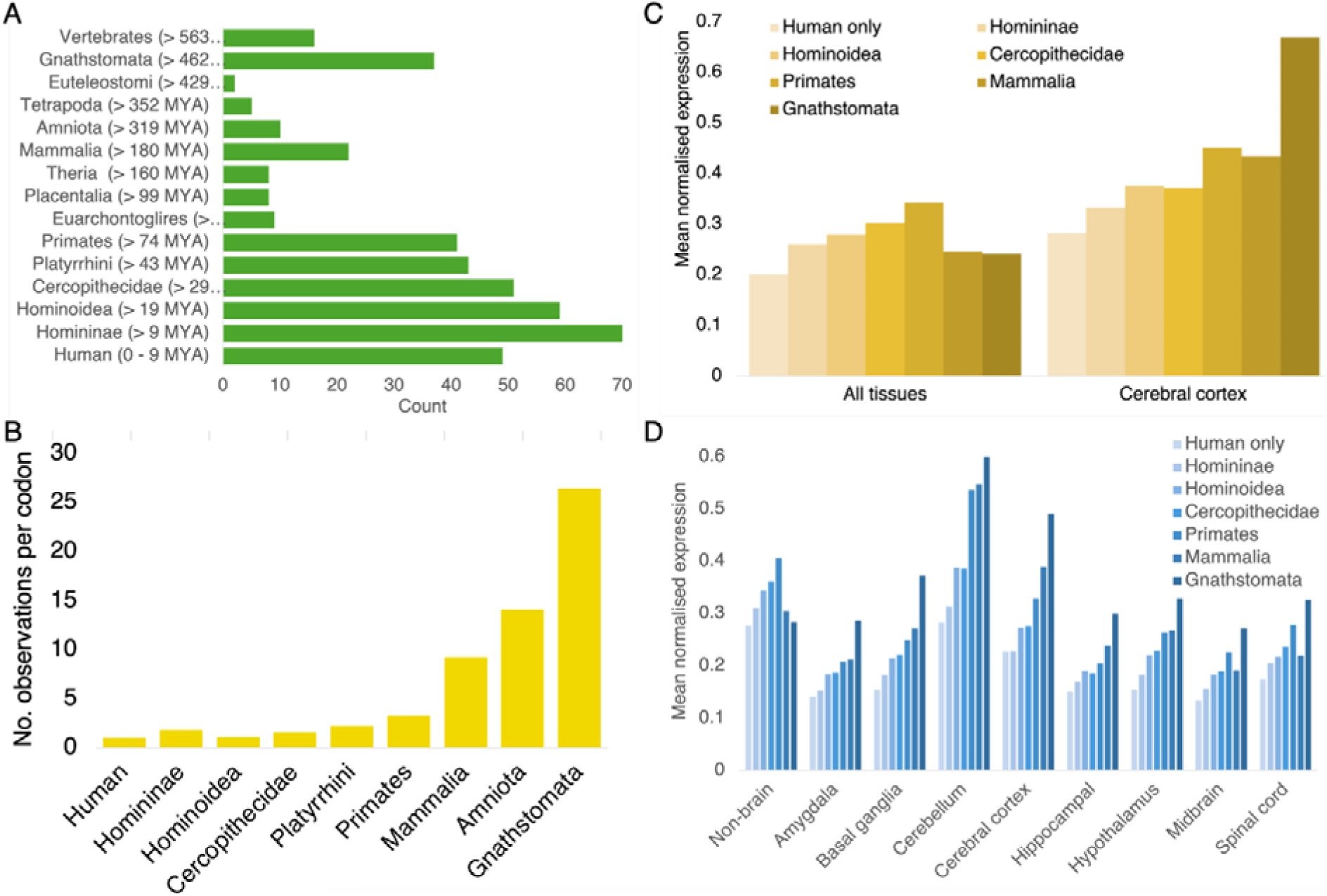
Dual coding region age and the quantity of supporting peptides. A. The age distribution of the 430 alternative splicing derived dual coding regions identified with PeptideAtlas peptide evidence. The y-axis shows the clade of the last common ancestor that preserves both frames and the estimation of the evolutionary distance of that last common ancestor taken from the TimeTree database (Kumar *et al*, 2020). B. The mean number of peptide observations mapping to the dual coding region per overlapping codon for each age group (we combined dual coding regions for some of the smaller clades), which is a rough estimate of dual coding isoform abundance. C. Mean normalised gene expression from the Human Protein Atlas large-scale tissue-based RNASeq analysis (Uhlen et al. 2015) for different age groups. To simplify the diagram, we combined dual coding regions for some of the clades and we show just the results for the cerebral cortex samples against the mean of all other tissues. D. Mean normalised gene expression from the GTEx large-scale tissue-based RNASeq analysis (GTEx Consortium, 2020) for each age group. To simplify, we combined dual coding regions for some age groups and we show the mean of the non-brain tissues, but include all the GTEX brain samples.

### The oldest dual coding regions have more peptide support

We grouped the dual coding regions by predicted age and calculated the peptide support (PeptideAtlas observations) for each group. The oldest dual frame coding exons had higher peptide support for their alternative reading frames. The 53 alternative isoforms that we estimated to have arisen at least 460 million years ago (those with alternative reading frames preserved across all jawed vertebrates) had an average of 460.4 PeptideAtlas observations per dual coding event, while the 272 alternative isoforms that were not even preserved within the whole primate clade had just 59 PeptideAtlas observations per dual coding event.

Older dual coding regions were also shorter, with a mean of 17.4 codons per event, compared to more than 40 for the unconserved dual coding regions. This also meant that the differences between age groups was even more accentuated when calculating the number of PeptideAtlas observations per overlapping codon (Figure 3B). The number of observations per codon gives a rough approximation of the abundance of the alternative isoforms in each age group. The 271 dual coding regions with alternative reading frames that were not preserved across all primates had just 1.5 observations per overlapping codon, compared to the 26.5 observations per codon for the 53 dual coding regions that were preserved across all jawed vertebrate species. The alternative isoforms produced from the oldest dual coding regions were almost 20 times more abundant.

### Genes with ancient dual coding regions are predominantly brain expressed

We investigated the relationship between the estimated age of the dual coding regions and tissue expression. We grouped the 430 genes with dual coding regions annotated in GENCODE v45 and supported by PeptideAtlas peptides by dual coding region age, though we grouped together several of the age groups because they had relatively few genes. For each group we calculated the mean normalised tissue expression from the expression data in the Human Protein Atlas (Uhlen *et al*, 2015). We used all three large-scale tissue-based expression analyses (see methods), the results for the Human Protein Atlas analysis can be seen in Figure 3C, the equivalent results generated from the GTEx (GTEx Consortium, 2020) can be seen in Figure 3D. Results from all three analyses showed that the genes with the older dual coding regions, in particular those conserved across all jawed vertebrates (Gnasthomata in Figure 3) were enriched in brain tissues, in cerebral cortex in Human Protein Atlas (Figure 3C), but in the multiple brain tissues analysed in both the GTEx and FANTOM analyses, but in particular in cortex, cerebellum and basal ganglia (Figure 3D).

For the Human Protein Atlas analysis, we carried out Mann-Whitney tests between the normalised expression levels in the cerebral cortex of genes with the oldest dual coding regions (those that could be traced back to jawed vertebrates) and their normalised expression in other tissues. Expression in the cerebral cortex was significantly higher than all other tissues among those genes. For example, compared to testis (the tissue with the second highest expression levels) the p-value was 0.00001 over the 51 genes. Expression in the cerebral cortex in the GTEx analysis was also significantly higher in the 64 genes with dual coding regions that were preserved at least across eutherian mammals (p = 4×10^−6^).

The clear signal from all large-scale analyses is that the older dual coding regions, and particularly those preserved across all jawed vertebrates, are expressed more frequently in brain tissues. A heatmap of the normalised expression from the FANTOM analysis for the genes with dual coding regions that are preserved across all jawed vertebrates illustrates this enrichment in expression in nervous tissues and in cerebral cortex, cerebellum and basal ganglia in particular (Figure 4).

**Figure 4.**
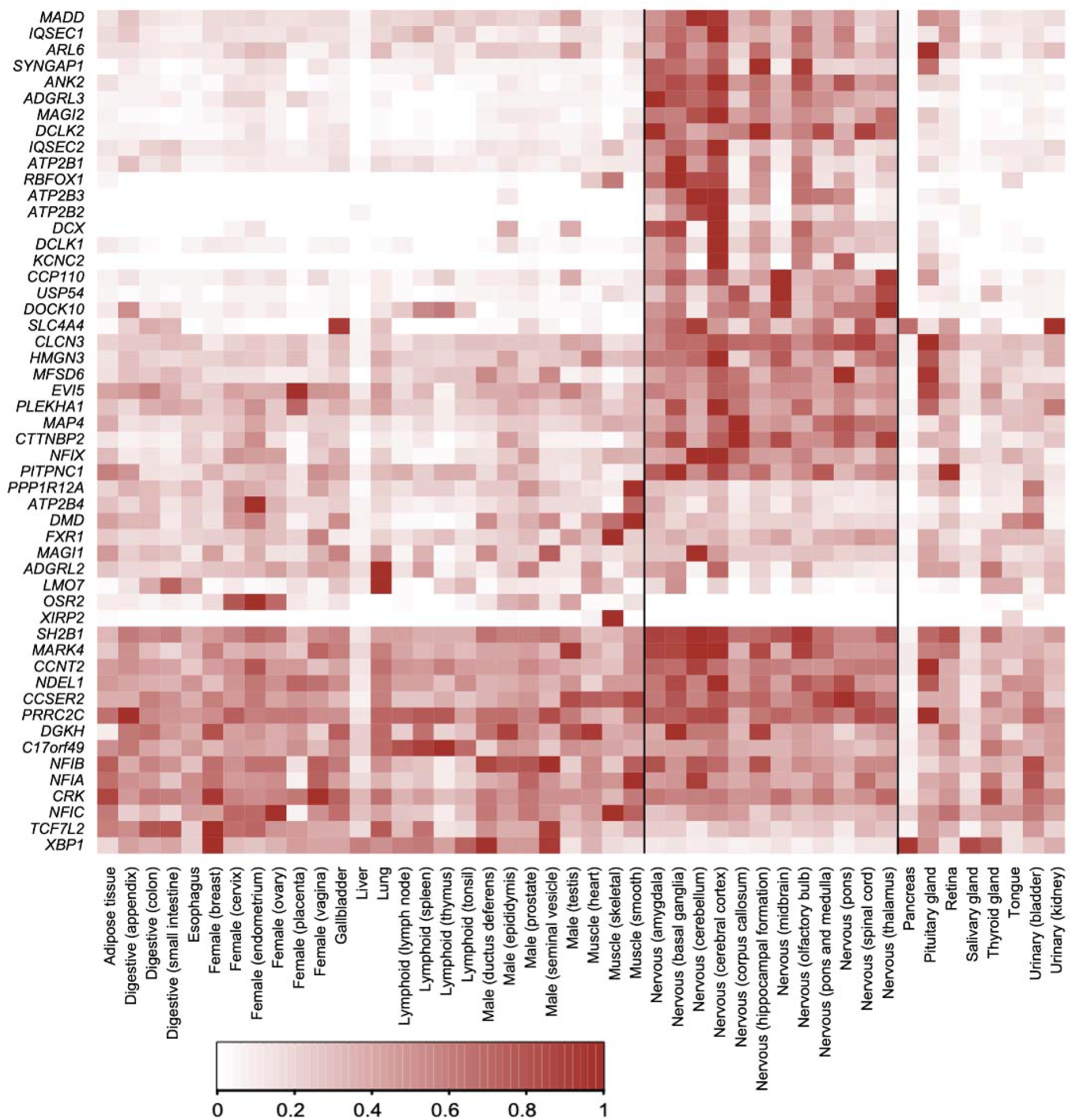
Tissue expression of jawed vertebrate dual coding regions. A heatmap of the normalised RNA expression of the 51 genes with dual coding regions that are conserved beyond the ancestor of jawed vertebrates over the 46 tissues from the large-scale tissue-based FANTOM analysis (Abugessaisa et al, 2017). The normalisation procedure is detailed in the methods section. Black vertical lines separate the nervous system tissues from other tissues.

### Close to 100 nonsense mediated decay targets are translated

Possibly the most unexpected result from the 430 dual coding regions with peptide evidence was that 21.9% had peptide support even though they should not have been translated. Alternative transcripts from 92 dual coding regions were predicted to be nonsense mediated decay (NMD) targets by annotators, and another two were tagged as non-stop decay targets. Yet all 94 of the alternative protein isoforms had peptide support in PeptideAtlas. All but 11 of the NMD targets came from dual coding regions that were not preserved beyond primate species.

We have detected peptide evidence for the translation of predicted NMD targets previously (Ezkurdia *et al*, 2012), but not at this scale.

### Manually annotated dual coding regions

To the 430 dual coding regions with PeptideAtlas support we added another 107 dual coding regions via manual curation. Sixty-five dual coding regions came from an analysis in which we mapped PeptideAtlas peptides to transcripts not annotated in GENCODE (Rodriguez *et al*, 2025). These alternative reading frames had peptide support but were only annotated in RefSeq or UniProtKB or the PeptideAtlas database itself (see methods).

A further 29 dual coding frames were annotated in the GENCODE v45 gene set but did not have peptide support for both frames. These were mostly paralogues (see methods). In many cases the reason that there was no peptide support was that one of the two C-terminals had no possible distinguishing tryptic peptides. These alternative reading frames all had conservation support. Finally, we found 13 novel alternative reading frames that did not have supporting peptides, but that were supported by deep conservation evidence (both reading frames were conserved at least to the ancestor of jawed vertebrates.

One unannotated alternative reading frame, in *ASXL1*, was supported by both peptides and deep conservation evidence. It has peptide support in PeptideAtlas because it was detected in a large-scale proteomics analysis (Ji *et al*, 2020) and it is conserved across all jawed vertebrates. Since there was no supporting evidence for alternative splicing, it had been postulated that the dual coding region in *ASXL1* and the equivalent dual coding region in its paralogue *ASXL2* arose from two separate conserved programmed ribosome profiling mechanisms (Dinan et al, 2017). The dual coding regions in the two paralogues appear to have little in common, but both do have a highly conserved HCF-1 binding motif in the alternative frame (Dinan et al, 2017). Despite the intriguing possibility that two distinct previously unknown and unrelated programmed ribosome profiling mechanisms occurred during the evolution of the two paralogues, the most parsimonious explanation for the evolution of the dual coding regions is still alternative splicing and gene duplication.

### Alternatively spliced dual coding regions and gene duplication

Across the extended set of 537 alternatively spliced dual coding regions, 90 were preserved across all jawed vertebrates. Many of these preserved alternative frames were found in paralogous genes, suggesting that the acquisition of dual coding exons predated the duplication of the paralogues. At least 13 gene families appear to have gained dual coding regions before gene duplication because these dual coding regions are clearly preserved in some or all of the members of the family. These families include the plasma membrane calcium-transporting ATPase 1 family (*ATP2B1*, *ATP2B2*, *ATP2B3* and *ATP2B4*, see Figure 5A), the nuclear factor 1 family (*NFIA*, *NFIB*, *NFIC* and *NFIX*), and the Ras/Rap GTPase-activating protein family (*SYNGAP1*, *RASAL2* and *DAB2IP*).

**Figure 5.**
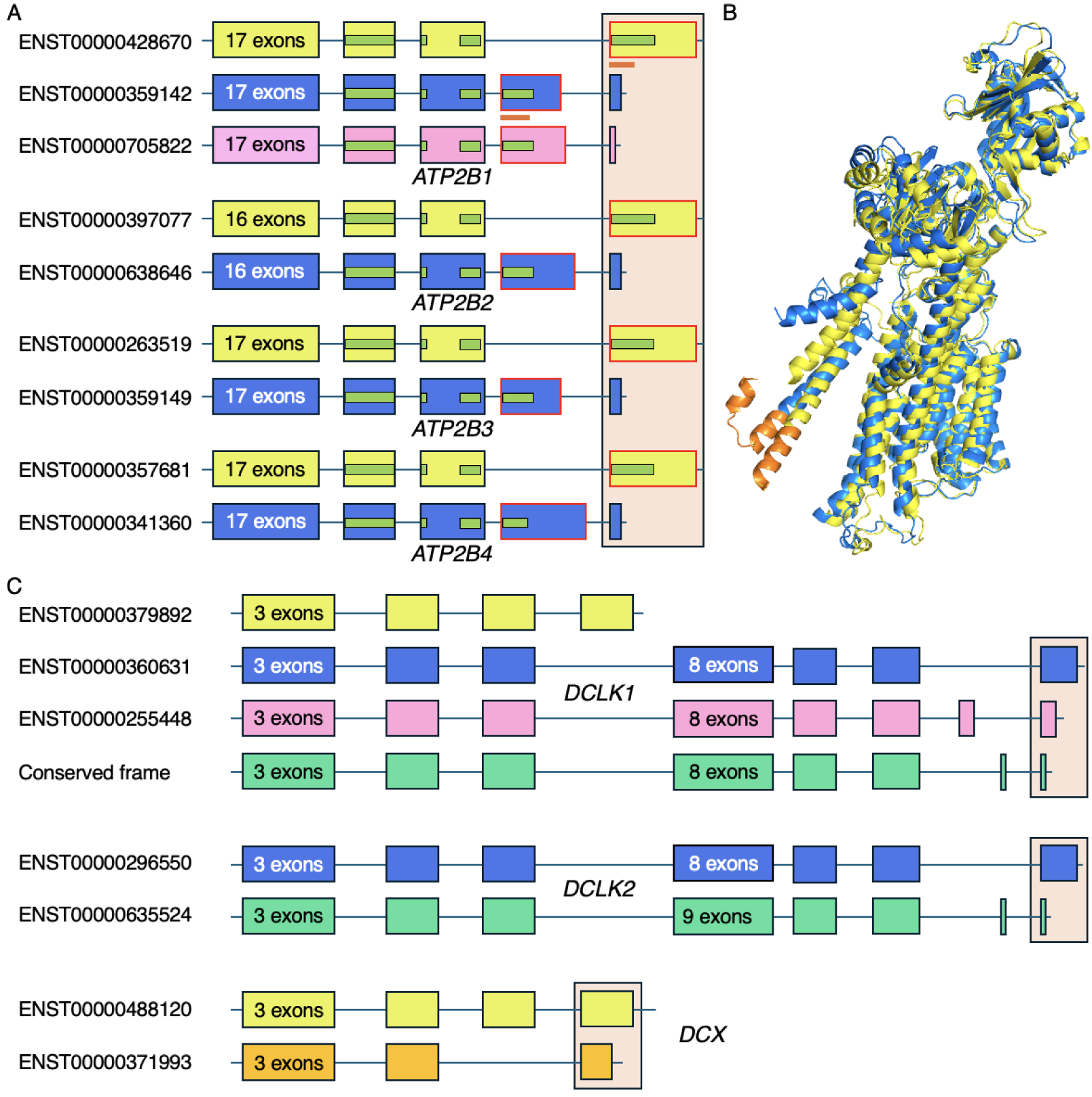
Two families with ancient dual coding regions. A. Dual coding regions in the 4 members of the plasma membrane calcium-transporting ATPase 1 family. Dual coding exons are highlighted by the light orange box. The ancestor of the plasma membrane calcium-transporting ATPase 1 family gained a C-terminal exon from a tandem duplication prior to the split between vertebrates and invertebrates (red outlined exons; Martinez Gomez *et al*, 2021) and the two duplicated C-terminals are used in a mutually exclusive fashion. In all vertebrate species the first duplicated exon gained a 3’ exon that overlaps the second tandem duplicated exon in a different frame. In *ATP2B1* this has happened twice, so there is a triple coding exon. This second dual coding region in transcript ENST00000705822 is preserved across all chordates. The smaller green rectangles show where Pfam (Mistry *et al*, 2021) domains map, the orange lines refer to the panel B structure. B. The AlphaFold (Abramson *et al*, 2024) predicted structures of two of the *ATP2B1* isoforms, those generated from ENST00000428670 (yellow) and ENST00000359142 (blue). The colours correspond to the exons in frame A. The only region that is different (corresponding to part of a Pfam domain) is shown in orange and corresponds to the short orange lines under the corresponding exons in panel A. Only a small fraction of the region that differs between ENST00000359142 and the other two isoforms is predicted to fold in a stable 3D structures by AlphaFold. C. The dual coding regions in the the doublecortin-like kinase family (*DCLK1* and *DCLK2*) and their paralogue, doublecortin (*DCX*). Dual coding exons are highlighted by the light orange box. One doublecortin-like kinase dual coding region is preserved across chordate species (the blue and green exons); the dual coding region unique to *DCLK1* and the dual coding region in *DCX* both have a last common ancestor with jawed vertebrates. There are indications that the short dual coding frame of *DCLK2* is preserved in *DCLK1* too, though the frame is not annotated as coding in *DCLK1*. If it is coding, then *DCLK1* also has a triple coding region.

Three gene families had dual coding regions that were preserved across chordate species. These were the Microtubule-associated protein family (*MAP4*, *MAP2* and *MAPT*), the MAP/microtubule affinity-regulating kinase family (*MARK1*, *MARK3* and *MARK4*), and the Doublecortin-like kinase family (*DCLK1* and *DCLK2*). As their name suggests, the MAP/microtubule affinity-regulating kinases phosphorylate the three Microtubule-associated proteins. Doublecortin-like kinases are also Microtubule-associated proteins kinases that bind at growing microtubule ends (Bechstedt and Brouhard, 2012), stimulate axon generation (Nawabi *et al*, 2015) and regulate dendritIc spines (Murphy *et al*, 2023). *DCX* (Neuronal migration protein doublecortin), a paralogue of *DCLK1* and *DCLK2* that does not have the C-terminal kinase domain, also has an ancient dual coding terminal exon (Figure 5C). The equivalent of the *DCX* dual coding exon is annotated in *DCLK1* but there is no indication that the second frame is preserved.

Seven alternatively spliced dual coding regions from three gene families that are preserved across all vertebrate species have PeptideAtlas peptide support for just one of the two reading frames. In each case this is because one of the two frames has few cleavable tryptic lysines or arginines. The three families are the Jun N-terminal kinase family (*MAPK8*, *MAPK9* and *MAPK10*), the pre-B-cell leukemia transcription factor family (*PBX1* and *PBX3*) and the muscleblind-like splicing regulator family (*MBNL1* and *MBNL2*).

### Conservation or protein support for 19 triple coding regions

Remarkably, there was either peptide or conservation evidence for 19 triple coding regions, exons that produce proteins from three distinct overlapping reading frames. Approximately half of the triple coding events are highly conserved and these overlapping regions are much shorter than the rest. In seven genes (*ADGRL2*, *ADGRL3*, *ATP2B1*, *CCSER2*, *DCLK1, MAP4* and *SYNCRIP*) all three reading frames are conserved across jawed vertebrate species. In another four genes, two of the three reading frames are conserved across jawed vertebrate species, *HMGN3* (two frames preserved across jawed vertebrates, one preserved across amniotes), *PRRC2C* (jawed vertebrates and eutheria), *RASAL2* (jawed vertebrates and mammals), and *SH2B1* (jawed vertebrates and euarchontoglires). Five of these 11 highly preserved triple coding frames, those in *ATP2B1*, *CCSER2*, *HMGN3*, *SH2B1* and *PRRC2C*, also have peptide support for all three frames.

### Comparison with previous genome-wide studies of dual coding regions

Between those with PeptideAtlas peptides and those with conserved alternative reading frames, the 537 dual coding regions from 490 genes that we found had some sort of functional evidence. We compared the genes that have dual coding regions that were annotated across the four studies (an analysis of the dual coding regions is not feasible). There was relatively little overlap between these genes and the genes with the dual coding regions from the previous genome-wide analyses (Figure 6). Just over a third of the 490 genes in our study coincided with the genes in the large Goubert set (Goubert *et al*, 2026), but fewer than 7% appeared in the earlier studies (Kovacs *et al*, 2010; Liang and Landweber, 2006). Almost two thirds (63.9%) of the genes in our analysis did not appear in any of the three previous genome-wide analyses. In fact, only 14 of the 1,699 genes across the four experiments appeared in all four analyses. This relatively minor overlap is not entirely surprising since none of the previous analyses based their selection of dual coding regions on proteomics evidence or cross-species conservation.

**Figure 6.**
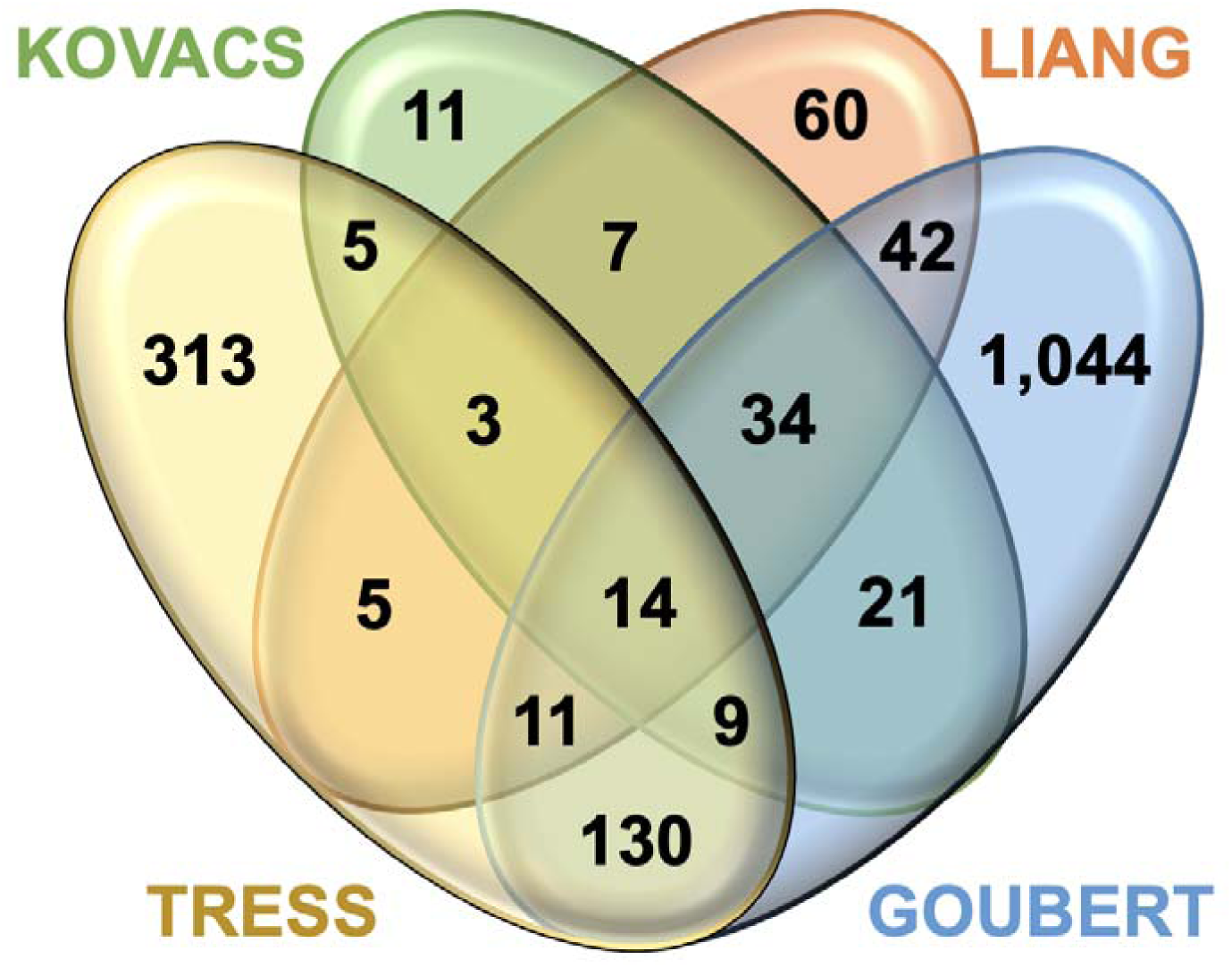
coincidence with previous studies. The overlap between the 490 genes with dual coding regions in our analysis (TRESS) and the genes reported to have dual coding regions in three previous large-scale analyses. The comparison is carried out at the gene level, if it were possible to compare by individual dual coding regions, the overlap between the four analyses would probably be even smaller.

### Half of all tissue specific protein isoforms are found in brain

Alternative splicing is often linked to tissue specificity (Glinos *et al* 2022) and Goubert *et al* (Goubert *et al*, 2026) carried out an analysis of transcript abundance for their dual coding regions using the GTEx experiments (GTEx Consortium, 2020). They noted a slight enrichment in brain specific transcripts. Their results suggest that this was largely due to a small number of genes that had ancient dual coding regions and brain specific transcripts, including *ATP2B1*, *ATP2B4*, *PBX1* and *MARK4* with dual coding regions that precede the earliest vertebrates, *CCSER2*, *MADD*, *MAGI1*, *PLCB4*, and *PTBP2* with dual coding regions that are preserved across all jawed vertebrates, and *GABBR1* that has a dual coding region that is preserved across all euteleostomi species. Eight of these dual coding regions were also in our analysis, but our manual curation did not find the *PTBP2* or *GABBR1* dual coding regions.

Instead of looking at tissue specificity at the transcript level, we searched for tissue specific protein isoforms produced from the 495 dual coding regions that had peptide support. In an earlier work we used proteomics data to investigate how widespread tissue specificity of alternative splicing was at the protein level (Rodriguez *et al*, 2020). We detected clear tissue specific alternative isoforms, and almost all the splice events that were tissue specific at the protein level were of ancient origin (Rodriguez *et al*, 2020).

Here we counted the number of times that isoform discriminating peptides were identified in tissue specific experiments (see methods section). Most experiments in PeptideAtlas involve dysregulated cells, cancer tissue, diseased tissue or cell lines, and these were ignored. After excluding these experiments, we found that most alternative isoforms in dual coding regions had too few tissue-based peptide identifications to determine whether the frame was tissue specific.

However, we did find evidence for tissue specific protein isoforms from 23 dual coding regions (Figure 7). Three dual coding regions had isoforms that were enriched in two different tissue groups, brain and immune tissues in the case of *CAMK1D*, and brain and heart in *ANK2* and *LMO7* dual coding regions (Figure 7). As we found in our earlier experiments (Rodriguez *et al*, 2020), most tissue specific alternative splice isoforms were of ancient origin; all but six of the 23 dual coding regions with tissue specific isoforms appeared to have evolved prior to the jawed vertebrate clade. Two dual coding regions were preserved only across old world monkeys.

**Figure 7.**
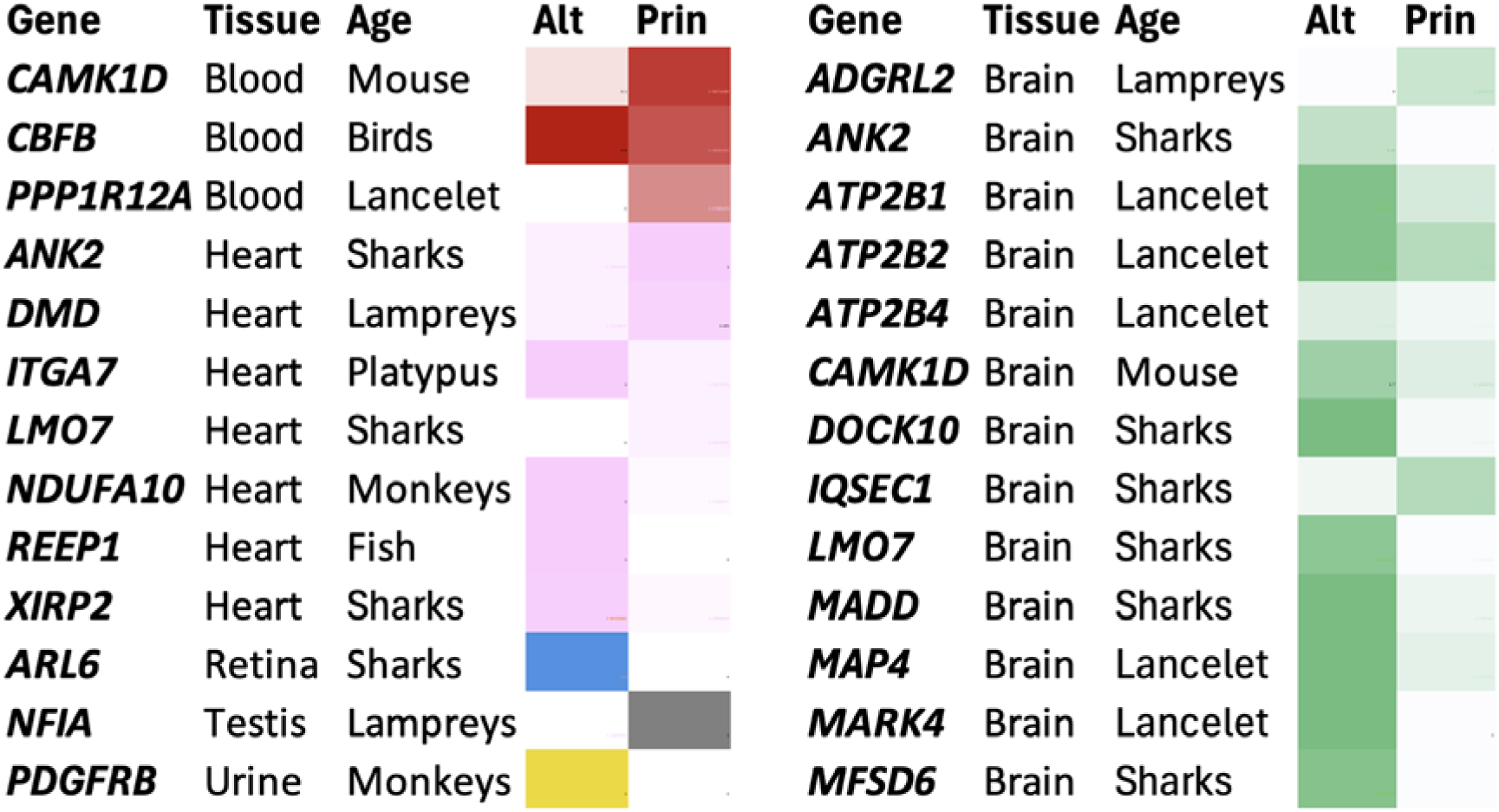
protein isoform tissue specificity. Heatmap of peptide support for dual coding frames in which either the alternative (Alt) or principal (Prin) coding frames were enriched in specific tissue groups. Heatmap colours correspond to distinct tissues, brain-enriched dual coding regions (green), blood-enriched (dark red), heart-enriched (pink), retina-enriched (blue), testis enriched (grey), urine-enriched (yellow). The column “age” indicates the oldest species in which the dual coding region is preserved. Most tissue-specific dual coding regions were already present in the last common ancestor of humans and sharks.

The protein level tissue specificity results reinforced the link between ancient dual coding regions and nervous tissues. Thirteen of the 23 dual coding regions produced nervous tissue enriched isoforms, while seven had isoforms that were enriched in heart and/or skeletal muscle. Reassuringly, four of the transcripts that had brain specific expression in the Goubert et al study (those in *ATP2B1*, *ATP2B4*, *MADD* and *MARK4*) had brain specific isoforms in our analysis. Between them, the two analyses suggest that genes with ancient dual coding regions are not only highly expressed in brain tissues (Figure 3), but that many ancient dual coding regions also produce brain specific protein isoforms.

For example, although we found that at the transcript level a large majority of those genes with ancient dual coding regions were highly expressed in the brain (Figure 4), a small number of the genes were not enriched in nervous tissues. Nevertheless, the two studies show that many of the genes that were not brain-enriched had brain-specific protein isoforms (*ADGRL2*, *ATP2B4*, and *LMO7*, Figure 7), or brain-specific transcripts (*CCSER2*, *MAGI1, PBX1*).

### Identifying alternative coding frames under selection pressure using dN/dS ratios

Analysis of cross species alignments showed that in all but a handful of the dual coding regions, one coding frame (the principal reading frame) is almost always under much clearer purifying selection than the other. This observation fits with the results of an analysis (Szklarczyk *et al*, 2007) that reported that the reading frames of most dual-coding exons were under unequal evolutionary pressures. The few exceptions to the rule we found were completely or almost completely conserved across their dual coding region.

We noted that not only was there stronger selection pressure in one reading frame, but also that alternative reading frames in the vast majority of dual coding regions had no clear evidence of any sort of selection constraints. Indeed, the reason that many alternative frames appear to be preserved may be related to the selection constraints that are imposed on the principal frames. Constraints on one frame of the dual coding region may mean that orthologous species have relatively few premature stop codons or frameshifts in the alternative reading frames.

Because one reading frame was clearly under greater selection pressure, we had to look beyond the overlapping regions to determine whether the alternative coding frames were under selection pressure too. The strategies we used are detailed in the methods section and described briefly here.

The first strategy was to evaluate the dN/dS of the alternative reading frame directly. This was possible only when the alternative reading frame extended beyond the principal reading frame, the frame that was under clear selective pressure (see Figure 2A). The second technique, which could be applied when the alternative reading frame was shorter, involved calculating dN/dS for any intervening exon or exon extension that changed the frame of the alternative exon (Figure 2B).

Finally, for those dual coding regions where the principal frame (under clear selection pressure) was longer than the alternative and there was no intervening frame changing exon, we divided the principal reading frame of the dual coding exon into two sections, the section that overlapped the alternative reading frame and the section that extended beyond the alternative reading frame (Figure 2C). We calculated dS scores for the principal reading frame of both sections. If amino acid changes are suppressed in the dual coding section, there ought to be substantially fewer synonymous mutations in related species and we would expect to find a lower dS for this section. We would expect the dS of the extended section ought to be considerably higher than that of the dual coding section because it is under selection pressure in just one frame. If the dS of the dual coding section is considerably lower than that of the extended section, this is an indication that both frames of the dual coding section are under selection pressure.

There were 35 dual coding regions that had dN/dS values that were significantly lower than 1 in two distinct reading frames of the same exon. This was clear evidence for selection pressure in both frames. This included the dual coding region in *DCLK1* (Figure 8), two that appeared to have evolved prior to the chordate clade (in *GULP1* and *MARK4*), and both dual coding frames from the Membrane-associated guanylate kinase, WW and PDZ domain-containing protein family (*MAGI1* and *MAGI2*). We also confirmed that the large unannotated dual coding regions in *ASXL1* and *ASXL2* were under selection pressure in two frames. All dual coding frames validated through dN/dS evaluation of the overlapping exon had dual reading frames whose evolution could be traced back prior to the eutherian clade, at least 99 million years ago.

**Figure 8.**
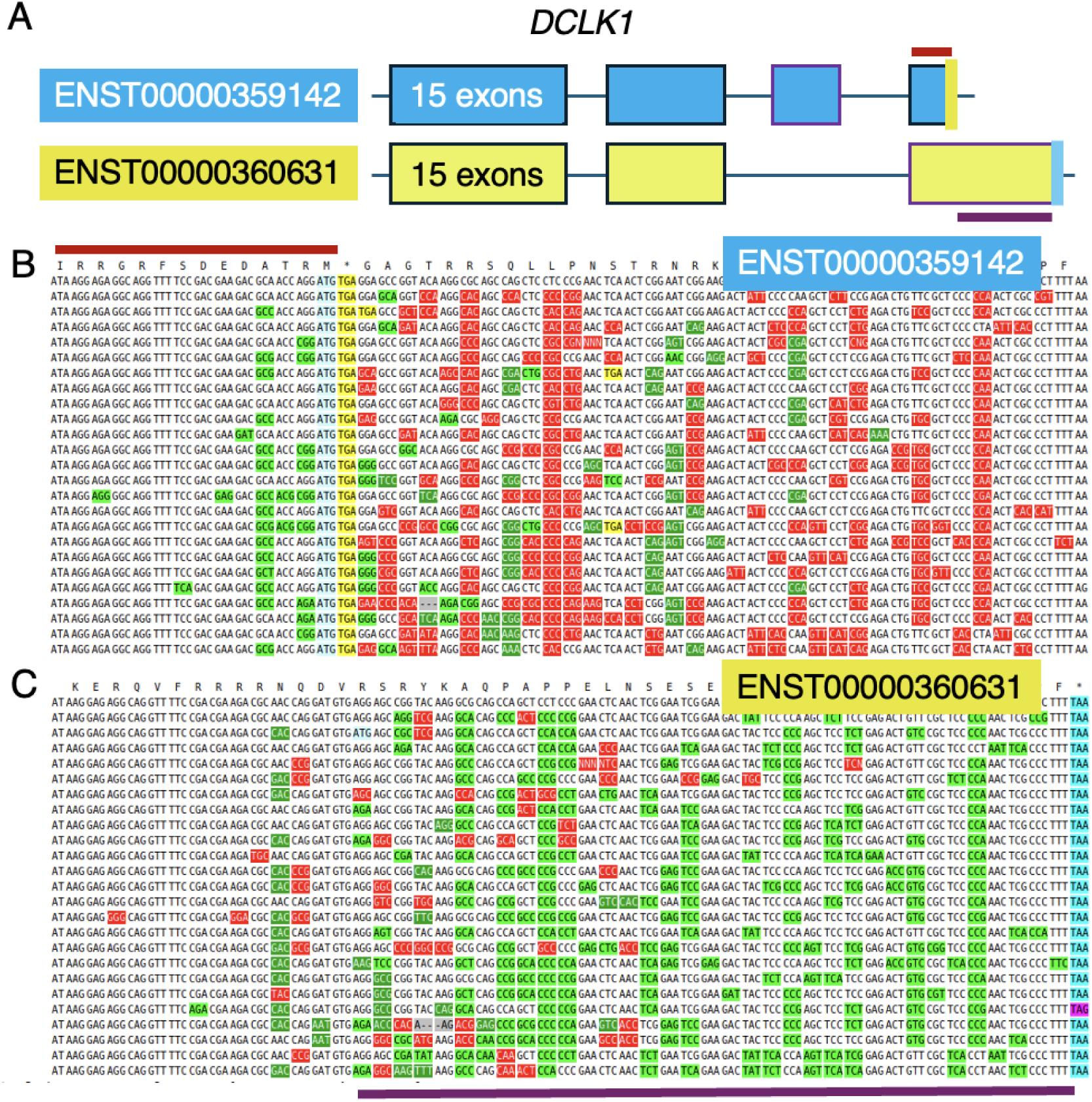
dN/dS analysis of two sections of the dual coding region in *DCLK1*. A. A representation of the two transcripts involved in the dual coding region annotated in *DCLK1*. The position of the red and purple lines corresponds to the image in panels B and C, the stop codons for the two reading frames are marked in yellow and blue and correspond to the stop codons in panels B and C. B. A representation of the cross-species alignment for ENST00000359142 over the region corresponding to the final coding exon of ENST00000360631 but read in a different reading frame; dN/dS was calculated over codons that are indicated by the red horizontal line for this reading frame. In panels B and C, codons are coloured by variant type, fluorescent green codons are synonymous with the human sequence, dark green and red codons are non-synonymous changes and stop codons are marked in yellow or blue (or purple). C. A representation of the cross-species alignment for the final coding exon of ENST00000360631; dN/dS was calculated over codons indicated by the horizontal purple line for this frame.

The dN/dS ratios for intervening frame changing exons and exon extensions (the second strategy) supported purifying selection in two different frames in another 18 dual coding regions. Examples validated using frame changing exons included the dual coding regions in *NFIA* and *NDEL1* (Figure 9). All 18 dual coding frames had alternative reading frames that evolved prior to the eutherian clade.

**Figure 9.**
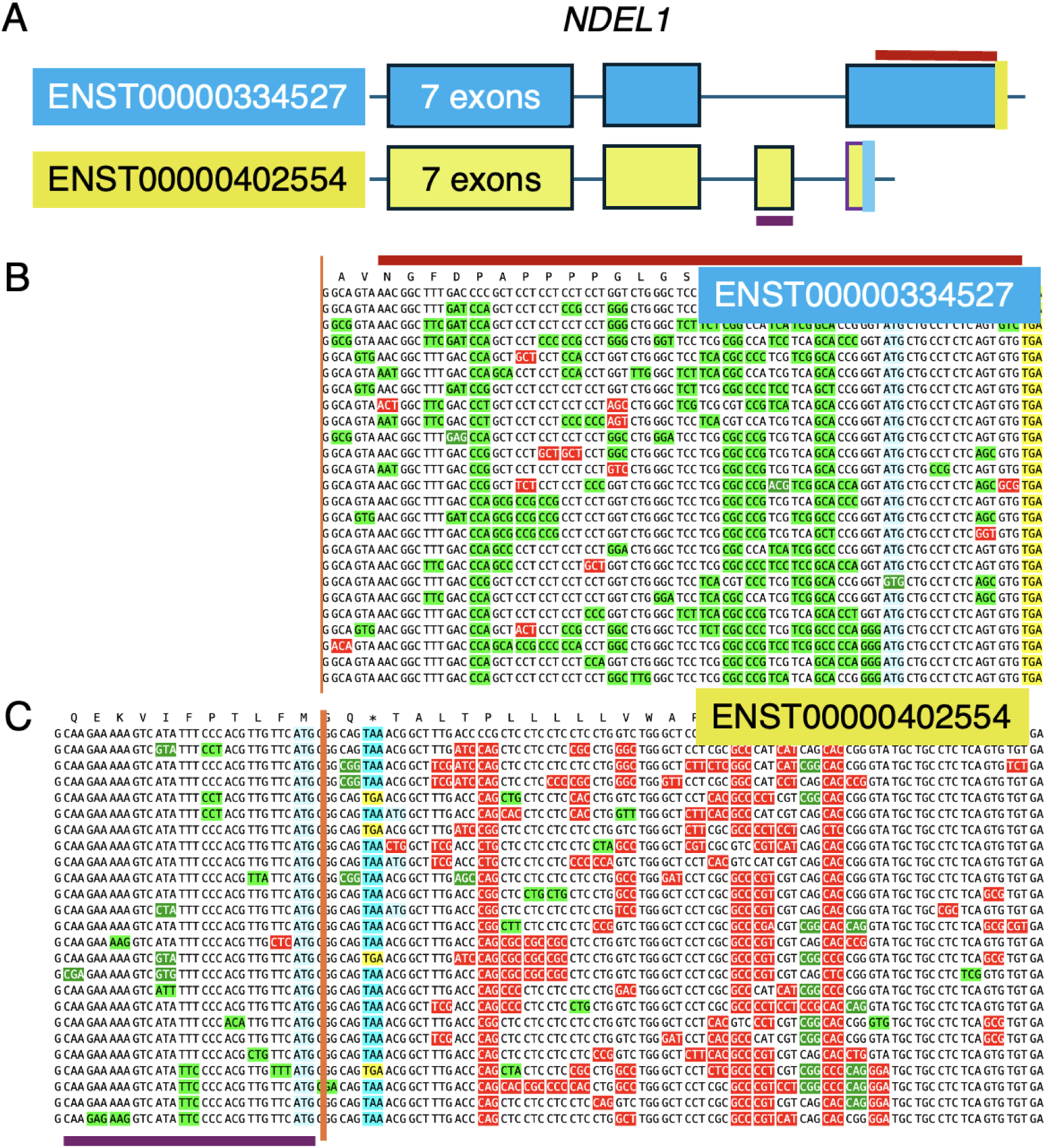
dN/dS analysis of dual coding region and frame changing exon in *NDEL1*. A. A representation of the two transcripts involved in the dual coding region in *NDEL1*. The position of the red and purple lines corresponds to the image in panels B and C, the stop codons for the two reading frames are marked in yellow and blue and correspond to the stop codons in panels B and C. B. A representation of the cross-species alignment for the final coding exon of ENST00000334527; dN/dS was calculated over codons shown by the horizontal red line for this frame. In panels B and C, codons are coloured by variant type, fluorescent green codons are synonymous with the human sequence, dark green and red codons are non-synonymous changes and stop codons are marked in yellow or blue. A representation of the cross-species alignment for ENST00000402554 combining the sequence of the upstream frameshifting exon with the region corresponding to the final coding exon of ENST00000334527 but read in a different reading frame; dN/dS was calculated over codons in the frameshifting exon, shown by the horizontal purple line, the vertical orange line marks the exon boundary.

### Using dS/dS ratios to detect dual coding regions under selection constraints

For the calculation of the dS/dS ratio based on the reading frame under most selection pressure (the principal reading frame, see Methods) we first calculated ratios for two groups of dual coding exons, a positive and a negative set. The positive set were 20 dual coding exons in which we had already shown that both reading frames had dN/dS ratios significantly lower than 1. We know that these dual coding regions are under selection pressure. The negative set was made up of 20 dual coding regions that were clearly under selection pressure in just one frame because closely related species had stop codons in the other reading frame. For the principal reading frame of all forty dual coding exons in the control set we plotted the dS values of the section of the exon that was predicted to be dual coding against the dS values of the extended section of the exon (predicted to code from just one reading frame).

Assuming a similar mutation rate, if an alternative frame in a dual coding region is under selection pressure, the dS values of the principal reading frame should be substantially lower in the dual coding section than in the extended (single coding) sections. The dS scores of the two control groups are plotted in Figure 10A. All 20 negative control dual coding regions had dS/dS ratios below 2.5, while all positive control dual coding regions had dS/dS ratios above 2.5 (Figure 10A).

**Figure 10.**
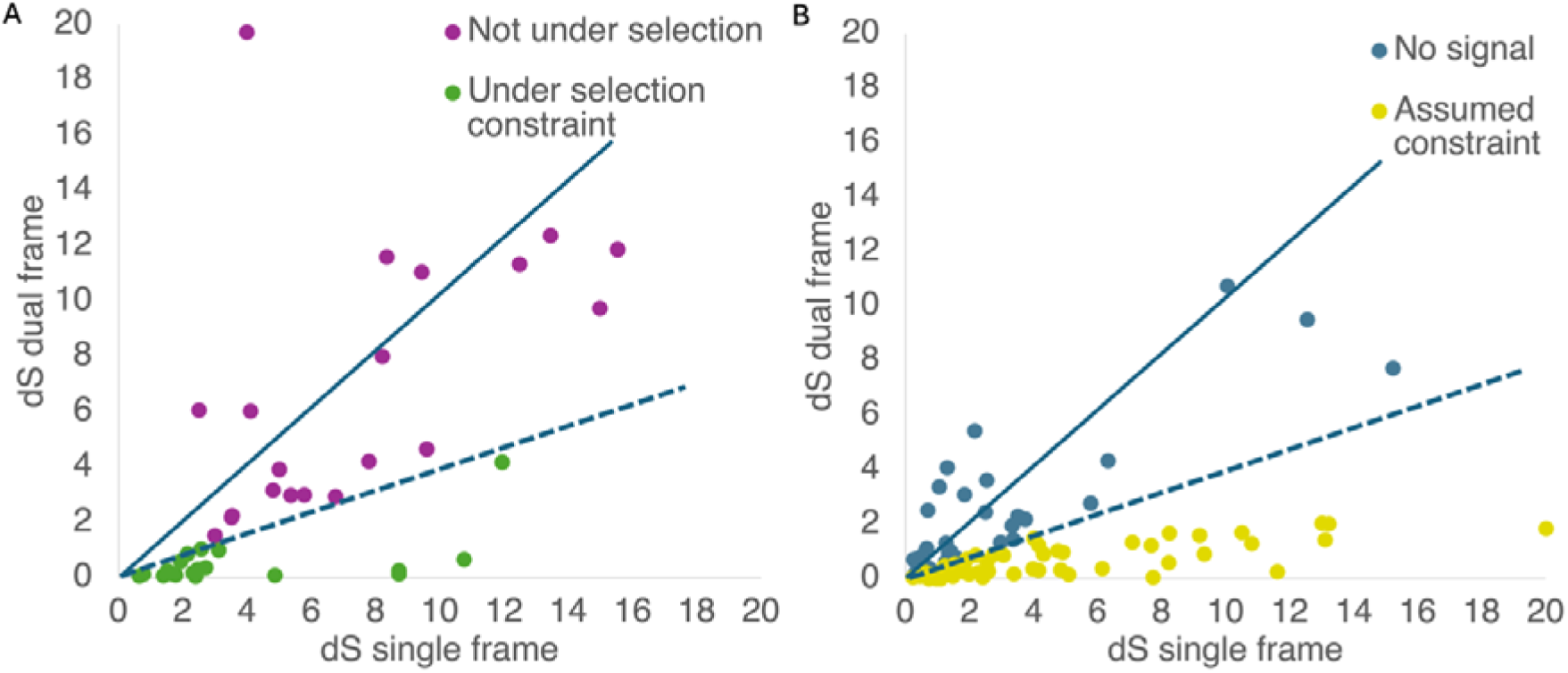
dS scores and the age of dual coding regions under selection pressure. A. Plot of control dS scores for different sections of each dual coding exon, on the x-axis the dS scores for the sections that code from a single reading frame, on the y-axis the dS scores for the sections that can code from two different reading frames. dS scores for the negative control set are in purple, dS scores for the positive control set are in green. The results for the negative control set, in which there is no selection in the alternative frame, would be expected to fall close to the full line. The dashed line separates the points of the two control sets. B. Plot of dS scores for tested dual coding exons, on the x-axis the dS scores for the sections that code from a single reading frame, on the y-axis the dS scores for the sections that can code from two different reading frames. dS scores above the cut-off are below the dashed line and highlighted in yellow, dS scores below the cut-off are above the dashed line in blue. The full line indicates where we would expect to find points from dual coding regions if only one frame was under selection.

Based on the control set, we chose a dS/dS ratio of 2.5 or greater as indicative of selection on the alternative coding frame. With this cut-off, we found a further 52 dual coding regions with indications that both frames were under selection pressure (Figure 10B). Genes with dual coding exons with dS ratios above the cut-off included the three IQ motif and SEC7 domain-containing genes (*IQSEC1*, *IQSEC2* and *IQSEC3*) and five alternative frames that were not annotated in any reference gene set in genes *GLCCI1*, *ADGRL2*, *MAP4*, *MAP2* and *MAPT*. The dual coding region in *GLCCI1* was detected because it was annotated in UniProtKB and had peptides in PeptideAtlas. The dual coding regions in *ADGRL2, MAP4*, *MAP2* and *MAPT* had equivalent reading frames in paralogues that were annotated and conserved across vertebrates. There was also evidence of selection pressure in two more recent dual coding frames, the dual coding region in *GAK*, which is only preserved across primate species, and the dual coding region in *TRIM47*, which we tagged as only conserved across old world monkeys. Although the exon extension that is required to change the frame is lost in simians and more distant species, the *TRIM47* dual coding region itself is conserved across mammals.

In total, 105 dual coding regions had evidence to support the existence of purifying selection in both frames. In addition, there were dual coding regions that we could not analyse with any of the three strategies, either because the overlapping or non-overlapping sections of the dual coding exons were too short, or because the cross-species alignments had next to no variation. For example, we believe that the gene *SYNCRIP* may produce proteins from three overlapping reading frames, but we are unable to test whether any of the three were under selection pressure because the final coding exon is identical across all tetrapod species. Thirty of the predicted dual coding regions that were preserved at least across all amniote species were either too short or too identical to be evaluated via dN/dS or dS/dS rations. Nevertheless, more than half of the dual coding regions that were preserved across eutherian species (56.7%) had evidence for selection in both frames (Figure 11).

**Figure 11.**
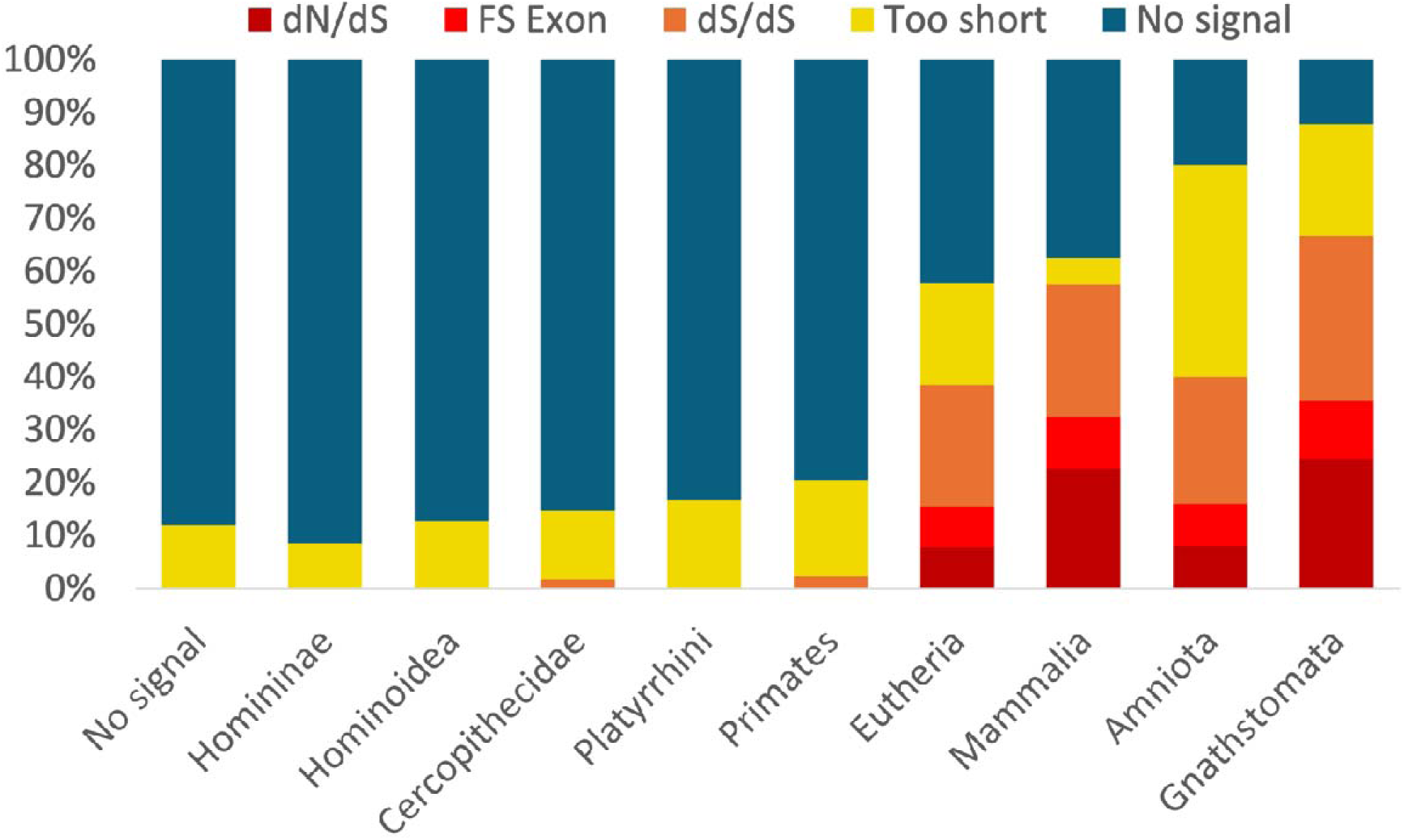
The relationship between selection constraints and dual coding region age. The proportion of dual coding regions with evidence for selection in two frames ordered by the minimum age of the dual coding region. “dN/dS” are those dual coding regions validated by the first strategy, “FS exon” are those dual coding regions validated by the second strategy, “dS/dS” are those dual coding regions validated by the dS analysis, “too short” are those dual coding regions that were either too short or too identical for the dS analysis.

### Multiple ancient dual coding regions are related to neurite outgrowth

Several of the ancient alternatively spliced dual coding regions in our set appear to have roles to the growth and differentiation of neurons. For example, the two *SHTN1* (shootin) isoforms that are produced from dual coding regions have different actin binding mechanisms (Ergin and Zheng, 2020), are expressed in different stages of axon growth and affect different aspects. The *Shtn1L* isoform is expressed first and promotes axon elongation by enhancing actin polymerization (Zhang *et al*, 2019). The *Shtn1S* isoform is produced later in the axon development process and promotes the specification of axons, determining axon fate (Kubo *et al*, 2015). The *SHTN1* dual coding region was traced back to a common ancestor that preceded the mammalian clade and both frames of the dual coding exon were validated in the dN/dS analysis.

We found two genes that had triple coding regions in which all three frames were highly conserved, were under detectable selection constraints and were supported by peptides. The first was *SH2B1*, which produces the SH2B adapter protein 1, a scaffold protein that promotes neurite outgrowth, the formation and branching of dendrites (Hsu *et al*, 2014; Chen *et al*, 2015) and the formation of the filopodia that precede the formation of axons and dendrites (Chen *et al*, 2015). The gene *SH2B1* can produce proteins with four different C-termini via an inserted exon, an exon extension and a 3’ exon with multiple coding frames (Figure 12A). These four isoforms are known as Sh2b1-alpha, -beta, -gamma and -delta (Yousaf *et al*, 2001).

**Figure 12.**
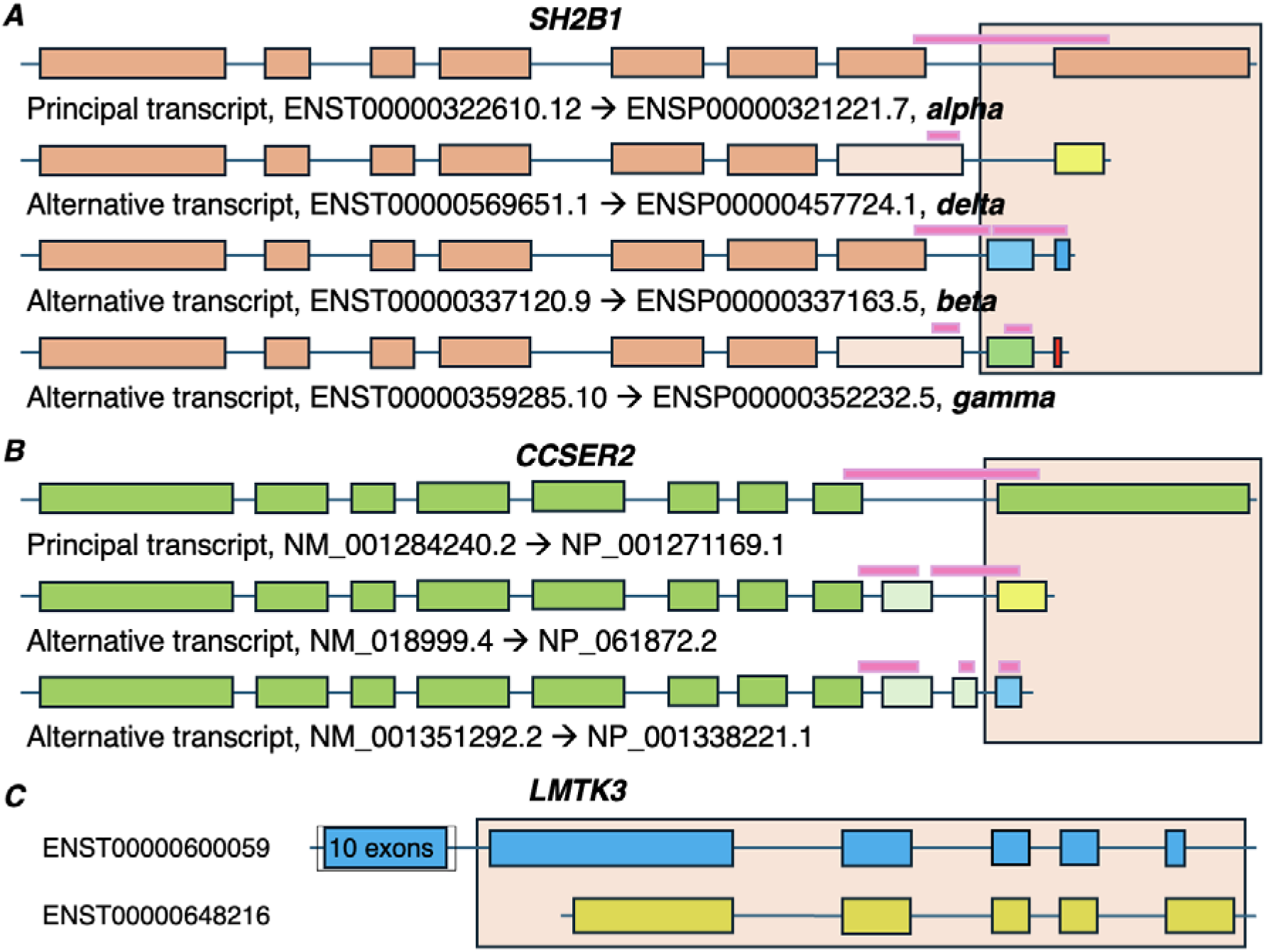
Dual coding regions in microtubule-related genes, *CCSER2*, *LMTK3* and *SH2B1*. A. The transcript models for the dual coding region in *SH2B1*: the four transcripts with different reading frames were all annotated in Ensembl/GENCODE so they are labelled with Ensembl identifiers along with the common names given to their isoforms (in Greek letters). The *SH2B1* dual coding region is marked by the light orange square and covers two exons. Exons in different frames are in different colours, and the pink horizontal bars show the peptides detected in PeptideAtlas experiments. B. The transcript models for the dual coding region in *CCSER2*: one of the three transcripts with a different reading frame is only annotated in RefSeq, so transcripts and isoforms are given RefSeq identifiers. The *CCSER2* dual coding region is marked by the light orange square and covers a single exon that is read in three different frames. Exons in different frames are in different colours, and the pink horizontal bars show the peptides detected in PeptideAtlas experiments. C. The transcript models for the dual coding region in *LMTK3*. The *LMTK3* dual coding region covers five exons and is highlighted with the light orange square. Exons in different frames are in different colours. Transcript ENST00000648216 is incomplete, it is missing the start codon. None of the exons in any of the panels are to scale.

The penultimate exon (used only in transcripts ENST00000337120 and ENST00000359285) codes from two different frames (marked in blue and green in Figure 12A), while the final exon can produce four different C-terminals from three coding frames. The -alpha and -gamma isoforms are produced from the same coding frame of the final exon, but in ENST00000359285 the final exon generates solely part of the stop codon. All four isoforms are important in neurite architecture and all increase neurite complexity, while the -delta isoform also increases neurite length (Cote *et al*, 2022). We find peptides to support all three frames and all three are under clear selection pressure.

The second gene with three frames under selection constraints and supported by peptides was *CCSER2* (see Figure 12B). The triple coding exon in *CCSER2* produces three different C-terminals that are all conserved at least as far as the earliest jawed vertebrates. The function of the different isoforms is not known, but the longest isoform is a tracking protein that is found at the end of extending microtubules (Shirai *et al*, 2023). Microtubules are the backbone of the developing neurites and their plus ends, closest to the growth cones, undergo variable phases of assembly and disassembly (Conde and Caceres, 2009; Prokop, 2013; Jakobs *et al*, 2022).

The main interactor of the *CCSER2* protein according to multiple experiments (Panneerselvam *et al*, 2024; Szklarczyk *et al*, 2023) is the nuclear distribution protein nudE-like 1 (*NDEL1*).

Interestingly, the *NDEL1* dual coding region is also conserved across jawed vertebrates and supported by peptides, and both frames are under purifying selection. *NDEL1* is necessary for the organization and anchoring of microtubules and plays an important role in the migration of newly formed neurons and regulates neurite outgrowth (Chansard *et al*, 2011).

The three families with the oldest dual coding regions, all with dual coding regions conserved to the ancestor of chordate species at least, also code for microtubule associated proteins and their proteins are either most expressed in nervous tissues or one of their isoforms is clearly brain specific. One of the three families is the doublecortin-like kinase family. The *DCLK1* kinase and its paralogue in *DCX* (another gene with an ancient dual coding exon) are microtubule-associated proteins involved in the regulation of neurite outgrowth and in particular the regulation of growth microtubules (Jean *et al*, 2012). Less is known about the second member of the family, *DCLK2*, but it has been shown that both *DCLK1* and *DCLK2* isoforms have multiple roles in the development of dendrites (Shin *et al*, 2013).

The other two families with ancient dual coding regions are related by function. *MAP2*, *MAP4*, and *MAPT* proteins all promote the polymerization of microtubules, and all affect the physical properties of microtubules in slightly different ways. These distinct properties contribute to the formation and differentiation of neurites (Nishida *et al*, 2023) *MARK1*, *MARK3* and *MARK4* produce kinases that specifically phosphorylate the *MAP2*, *MAP4*, and *MAPT* isoforms. The *MARK1* kinase has been shown to play a role in axon formation (Reiner and Sapir, 2014) and to regulate the development of dendritic spines (Kelly-Castro *et al*, 2024), and the *MARK4* kinase, which is localized prominently at the growing ends of neurite-like processes, is predicted to be directly involved in the organization of microtubules in developing neurons (Trinczek *et al*, 2004).

The *LMTK3* dual coding region is possibly the most intriguing since it spans five exons and a total of 573 codons (Figure 12C). Lemur tyrosine kinase 3 is a serine threonine kinase and almost entirely expressed in nervous tissues. It too has been implicated in the regulation of microtubule stability (Cilibrasi *et al*, 2021) and it has been suggested that it plays an important role in the arrangement of synaptic structure, synaptic plasticity and neurite outgrowth (Larose *et al*, 2024). As yet no work appears to have been carried out on the protein produced by the alternative frame.

We detected 49 tryptic peptides with a total of 1268 observations for the isoform coded from the alternative reading frame of *LMTK3* in PeptideAtlas. Peptides for both the principal and alternative frames were detected exclusively in brain tissues (both healthy and diseased). The alternative frame is preserved across mammalian species, although the alignments with marsupial and monotreme species are poor. Not only that, but both frames are under purifying selection based on dN/dS. The alternative frame of *LMTK3* is not annotated as a complete gene model in the human gene build. There is a full gene model of a transcript with the alternative frame in the Ensembl/GENCODE mouse gene set, but the conservation and peptide evidence we found suggests that the mouse model is not correct.

### Genes with ancient dual coding frames are Involved in neuron development

In total 82 genes had dual coding regions that were preserved at least to the ancestor of all jawed vertebrates. We analysed these genes with the online functional tool DAVID (Sherman *et al*, 2024) to find out whether the genes with these ancient alternative splice events were enriched in any particular functional processes. The top 10 groups of enriched GO terms are related to the formation of neuron projections and synapses (table 1).

**Table 1.**
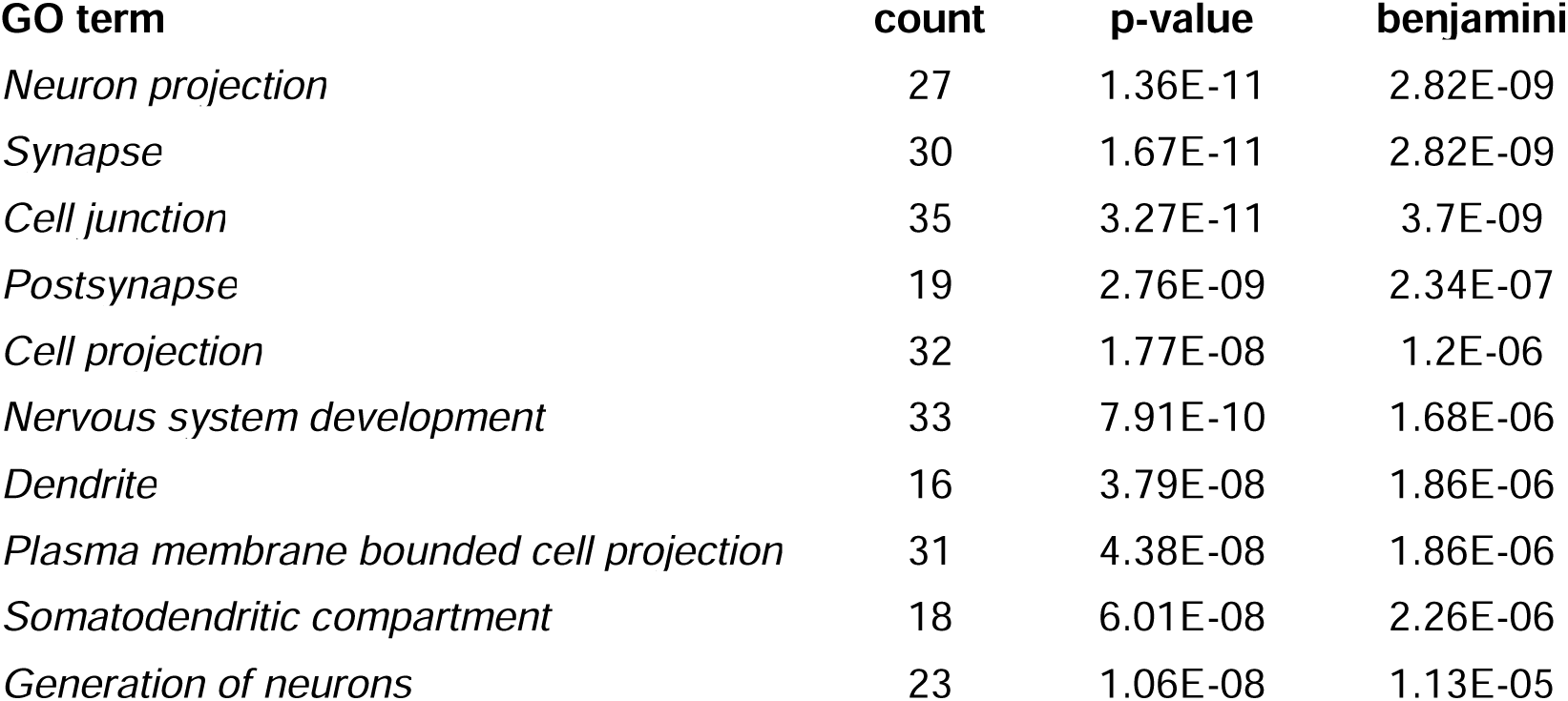
The top 10 Gene Ontology terms for dual coding regions conserved across jawed vertebrates. Gene Ontology terms for the 82 genes with dual coding regions preserved across all jawed vertebrates. GO terms were calculated using the GO Terms All option of the DAVID functional annotation system and sorted by Benjamini-Hochberg score.

Almost two thirds of the genes with dual coding regions preserved across jawed vertebrates are annotated with GO terms related to the generation of neurons, synapses and neural cell projections. In total, 49 of the 81 genes are annotated with two or more of these GO terms. Five genes, *ATP2B2*, *DOCK10*, *MARK1*, *MAPT* and *SYNGAP1*, coincide in all ten GO terms and three more, *NDEL1*, *KCNC2* and *FXR1*, appear in 9 of the top 10 GO terms.

A literature search with the other 33 genes found that another 18 are involved in the formation or regulation of neural cell projections, nervous cell development or synapses. These include *EVI5* and *BICD1* that have been found to interact to regulate abnormal dendrite branching in *C. elegans* (Fang *et al*, 2024), and *CCSER2* and *SH2B1*, the two genes with triple coding frames under clear selection pressure. Including these genes, we believe that at least 67 of the 82 genes with dual coding frames preserved across jawed vertebrates (81.7%) are implicated in neural cell projections, the development of neurons or the formation of synapses. Four of the remaining 15 genes, *DGKH*, *LMO7*, *MFSD6*, and *PITPNC1*, are known to be expressed specifically in neurons or to have neuron-specific transcripts. All 30 of the genes with the oldest dual coding regions, those that were preserved across vertebrate species, either had GO terms related to neuron development, differentiation and projection, or had literature to support their role in these processes.

## Conclusions

In this analysis we identified a set of 537 alternatively spliced dual coding regions that are either supported by peptide evidence or preserved at least across all mammalian species. In all dual coding regions the choice of reading frame is determined by an upstream alternative splicing event and the reading frames produce different protein isoforms, almost always with distinct C-terminals. Proteins with dissimilar C-terminals are likely to have diverse binding partners or posttranslational modifications, while changing the C-terminal may also affect protein stability (Sharma and Schiller, 2019; Chu *et al* 2026).

We based our initial investigation on the Ensembl/GENCODE human gene set, in which there are more than 6,000 coding genes that are annotated with dual coding regions. We found 430 dual coding regions with PeptideAtlas peptide support though more than 60% of these were not preserved beyond simians. These novel dual coding regions generally had little peptide support, and in most cases peptides were detected only in cancer cells or cell lines. It is possible that many of these dual coding regions are translated only as a result of aberrant splicing. The link between aberrant splicing and cancer is well established (Fackenthal and Godley, 2008; Inoue and Fry, 2016; Cherry and Lynch, 2020).

However, among the 430 Ensembl/GENCODE dual coding regions with peptide support we did find more than 100 apparently highly conserved dual coding regions, including 53 that were preserved across all jawed vertebrate species. Alternative isoforms produced from these dual coding regions were the most abundant.

One surprising result was that more than a fifth of all the alternative transcripts that produced peptides were predicted to be NMD targets. These transcripts should not have been translated and the fact that we found so much evidence for apparent NMD targets suggests either that either NMD is a highly noisy biological process, or that the rules for defining NMD transcripts need to be revisited. In their analysis, Goubert *et al* (Goubert *et al*, 2026) found that more than a third of the alternative isoforms listed in the UniProtKB database would likely be products of NMD and theorized that many of these transcripts might have roles related to gene expression rather than produce protein isoforms. We find that many of these transcripts are translated.

We added 107 ancient dual coding regions that we found by manual curation, including 78 dual coding regions that were not annotated by Ensembl/GENCODE. This brought the total of dual coding regions preserved across all eutherian mammals to 169. Ninety of these appeared to predate the ancestor of jawed vertebrates so would be more than 460 million years old and therefore these dual coding regions would be some of the oldest known alternative splicing events (Martinez Gomez *et al*, 2022). Remarkably, 19 genes had evidence to support translation from three coding frames, and twelve of these were preserved at least across all mammalian species.

We used dN/dS and dS/dS ratios to determine whether there was evidence for selection pressure on both frames among dual coding regions. We found 105 dual coding regions with evidence to suggest selective constraints in both reading frames. Genes *CCSER2* and *SH2B1*, with roles in directing microtubules (Shirai *et al*, 2023) and filopodia (Chen *et al*, 2015) in neurite development, had significantly low dN/dS ratios in all three frames. The number of dual coding regions we predict to be under functional constraints is likely to be an underestimate, because 29 dual coding regions that have been preserved across mammalian species were either too short or had too few mutations to be analysed with any of the strategies that we used for detecting selection constraints.

Many groups have found a relationship between alternative splicing events and nervous tissues (Grabowski, 2011; Li *et al*, 20156; Vuong *et al*, 2016; Zheng, 2020). What is remarkable in our analysis is the proportion of genes with ancient dual coding regions that are brain enriched and in particular have roles related to neuron development.

We find that genes with ancient dual coding regions are highly expressed in nervous tissues, especially in cerebral cortex and cerebellum, and further support for a relationship between ancient alternatively spliced dual coding regions and brain-specific function comes from the evidence for brain specific splicing at the protein level, at the transcript level (Goubert *et al*, 2026) and even among genes that are less expressed in brain tissues.

We find that genes with ancient dual coding regions are extremely enriched in functional terms related to neuron development. A literature search confirmed that 67 of the 82 genes with house dual coding regions that can be traced back to the ancestor of jawed vertebrates had functional roles related to the formation of neurons, dendrites or synapses, or to nervous system development in general. This included all 34 genes with dual coding regions that predated the evolution of the first vertebrates.

Taken together our evidence strongly suggests that ancient alternatively spliced dual coding regions play important roles in the formation and regulation of developing neurons and the formation of synapses, and that these dual coding regions may have been highly influential in the evolution of the vertebrate central nervous system.

## Acknowledgments

No large language models, chatbots, or any other form of artificial intelligence were used in the collection, curation, or analysis of data in this document, nor in the preparation and writing of the text. This work is 100% human.

